# Post-Transplant Administration of G-CSF Impedes Engraftment of Gene Edited Human Hematopoietic Stem Cells by Exacerbating the p53-Mediated DNA Damage Response

**DOI:** 10.1101/2023.06.29.547089

**Authors:** Daisuke Araki, Vicky Chen, Neelam Redekar, Christi Salisbury-Ruf, Yan Luo, Poching Liu, Yuesheng Li, Richard H. Smith, Pradeep Dagur, Christian Combs, Andre Larochelle

**Affiliations:** Cellular and Molecular Therapeutics Branch, National Heart, Lung and Blood Institute (NHLBI), National Institutes of Health (NIH), Bethesda, MD 20892, USA; Integrated Data Science Services (IDSS), National Institutes of Allergy and Infectious Diseases (NIAID), NIH, Bethesda, MD 20892, USA; DNA Sequencing and Genomics Core Facility, NHLBI, NIH, Bethesda, MD 20892, USA; Flow Cytometry Core Facility, NHLBI, NIH, Bethesda, MD 20892, USA; Light Microscopy Core Facility, NHLBI, NIH, Bethesda, MD 20892, USA

## Abstract

Granulocyte colony stimulating factor (G-CSF) is commonly used as adjunct treatment to hasten recovery from neutropenia following chemotherapy and autologous transplantation of hematopoietic stem and progenitor cells (HSPCs) for malignant disorders. However, the utility of G-CSF administration after *ex vivo* gene therapy procedures targeting human HSPCs has not been thoroughly evaluated. Here, we provide evidence that post-transplant administration of G-CSF impedes engraftment of CRISPR-Cas9 gene edited human HSPCs in xenograft models. G-CSF acts by exacerbating the p53-mediated DNA damage response triggered by Cas9- mediated DNA double-stranded breaks. Transient p53 inhibition in culture attenuates the negative impact of G-CSF on gene edited HSPC function. In contrast, post-transplant administration of G-CSF does not impair the repopulating properties of unmanipulated human HSPCs or HSPCs genetically engineered by transduction with lentiviral vectors. The potential for post-transplant G-CSF administration to aggravate HSPC toxicity associated with CRISPR-Cas9 gene editing should be considered in the design of *ex vivo* autologous HSPC gene editing clinical trials.

**Graphical Abstract:** 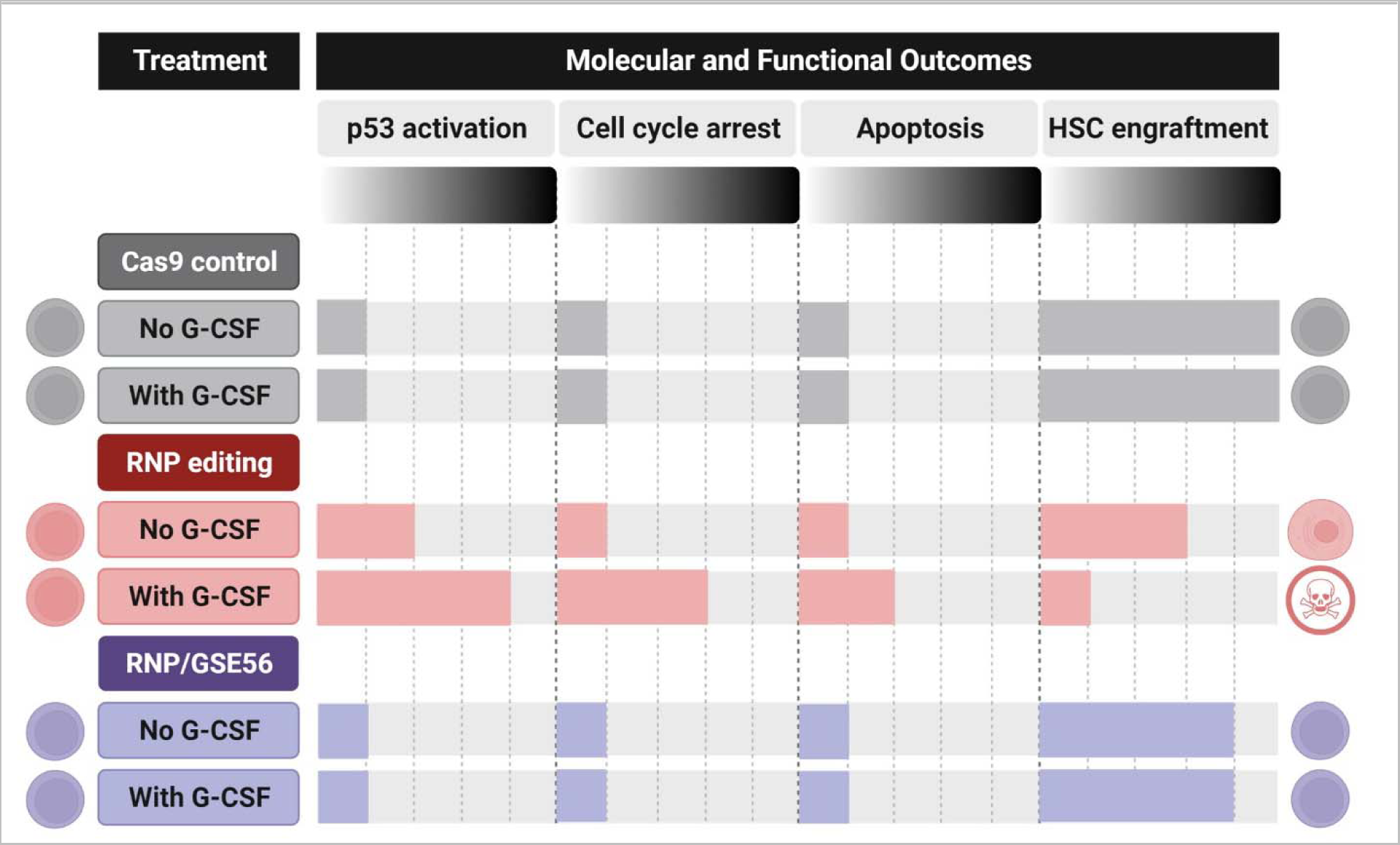

## Main

In recent years, transplantation of genetically modified autologous hematopoietic stem and progenitor cells (HSPCs) has provided clinical benefit in subjects with inherited hematological and immunological disorders. Transfer of a therapeutic gene to HSPCs has been successfully achieved using replication-defective integrating lentiviral vectors^1–6^ and, more recently, the programmable CRISPR-Cas9 nuclease system for precise genome editing^7–9^. The RNA-guided (gRNA) Cas9 nuclease induces site-specific DNA double-stranded breaks (DSBs) within the genome of target HSPCs, triggering transient activation of p53-mediated DNA damage response (DDR) and cell cycle arrest to enable DNA repair. In the context of Cas9- mediated homology directed repair (HDR), the presence of an HDR DNA donor template delivered via recombinant adeno-associated virus serotype 6 (AAV-6) results in an even more robust p53 transcriptional response, suggesting a cumulative p53 pathway activation from converging inputs^10–12^. These cellular responses impaired HSPCs’ ability to reconstitute and maintain hematopoiesis long-term in recipient hosts^10–15^. Hence, strategies to mitigate HSPC toxicity mediated by DNA damaging programmable nucleases have significant clinical relevance in *ex vivo* HSPC gene therapy applications.

The hematopoietic cytokine granulocyte-colony stimulating factor (G-CSF) and its cognate receptor (G-CSFR) are the foremost regulators of granulopoiesis. G-CSF plays an essential role in promoting granulocytic output by accelerating maturation of marrow progenitors into mature neutrophils and by facilitating their release into the circulation^16^. Clinical guidelines recommend administration of G-CSF after chemotherapy and autologous transplantation for hematologic malignancies to reduce duration of severe neutropenia and the associated risks of life-threatening infectious complications^17^. However, the potential benefits and shortcomings of G-CSF as an adjunct therapy following transplantation of gene corrected autologous HSPCs for inherited blood disorders has not been evaluated. In most studies, clinical practices have been extrapolated from guidelines recommended based on transplantation of unmanipulated autologous HSPCs.

The G-CSF:G-CSFR signaling axis is also active in multipotent hematopoietic stem cells (HSCs) with long-term (LT) repopulating potential (LT-HSCs) and genetically engineered G-CSFR^-/-^ mice have impaired HSC function^18, 19^. In experimental models, G-CSF was shown to potentiate DDR elicited by physiologic stressors, such as systemic infections or chronic blood loss, and by exogenous administration of chemotherapeutic agents or ionizing radiation^20–24^. Addition of G-CSF constricted the primitive hematopoietic compartment by promoting differentiation and/or senescence of HSCs at the expense of their self-renewal capacity^25–28^. Here, we hypothesized that post-transplant administration of G-CSF might also negatively impact HSCs treated with DNA-damaging programmable CRISPR-Cas9 nucleases. Our data indicate that G-CSF reduces the long-term repopulating potential of CRISPR-Cas9 gene edited human HSPCs after transplantation, exposing a previously unexplored facet of *ex vivo* autologous HSPC gene editing for the treatment of human diseases.

## Results

### Post-transplant administration of G-CSF impedes long-term engraftment of CRISPR-Cas9 gene edited HSPCs

As a proof-of-principle, we evaluated the impact of G-CSF use post-transplantation on the repopulating function of human HSPCs genetically edited by CRISPR-Cas9-mediated insertion/deletion (indel) formation at the AAVS1 “safe harbor” genomic locus^29^. Human mobilized peripheral blood (MPB) CD34+ cells from a healthy donor were pre-stimulated for 2 days and subsequently electroporated with either AAVS1-specific sgRNA/Cas9 ribonucleoprotein (RNP) complexes (designated “RNP group”) or Cas9 alone (designated “Cas9 group”). In the RNP group, we confirmed an editing efficiency of 61% in bulk CD34+ HSPCs, as determined by T7 endonuclease I (T7EI) cleavage of target site PCR amplicons (Extended Data Fig. 1a). These CRISPR-Cas9 edited cells were subsequently transplanted into immunodeficient NOD-scid lL2Rg^null^ (NSG) mice conditioned with a single non-myeloablative dose of busulfan (25 μg/kg), as commonly used in clinical HSPC gene therapy protocols^30^. We subcutaneously injected G-CSF (filgrastim, 125 μg/kg/day) or PBS (designated “untreated group”) from post-transplant day 1 to 14 and compared hematopoietic reconstitution (Fig. 1a).

**Fig. 1.**
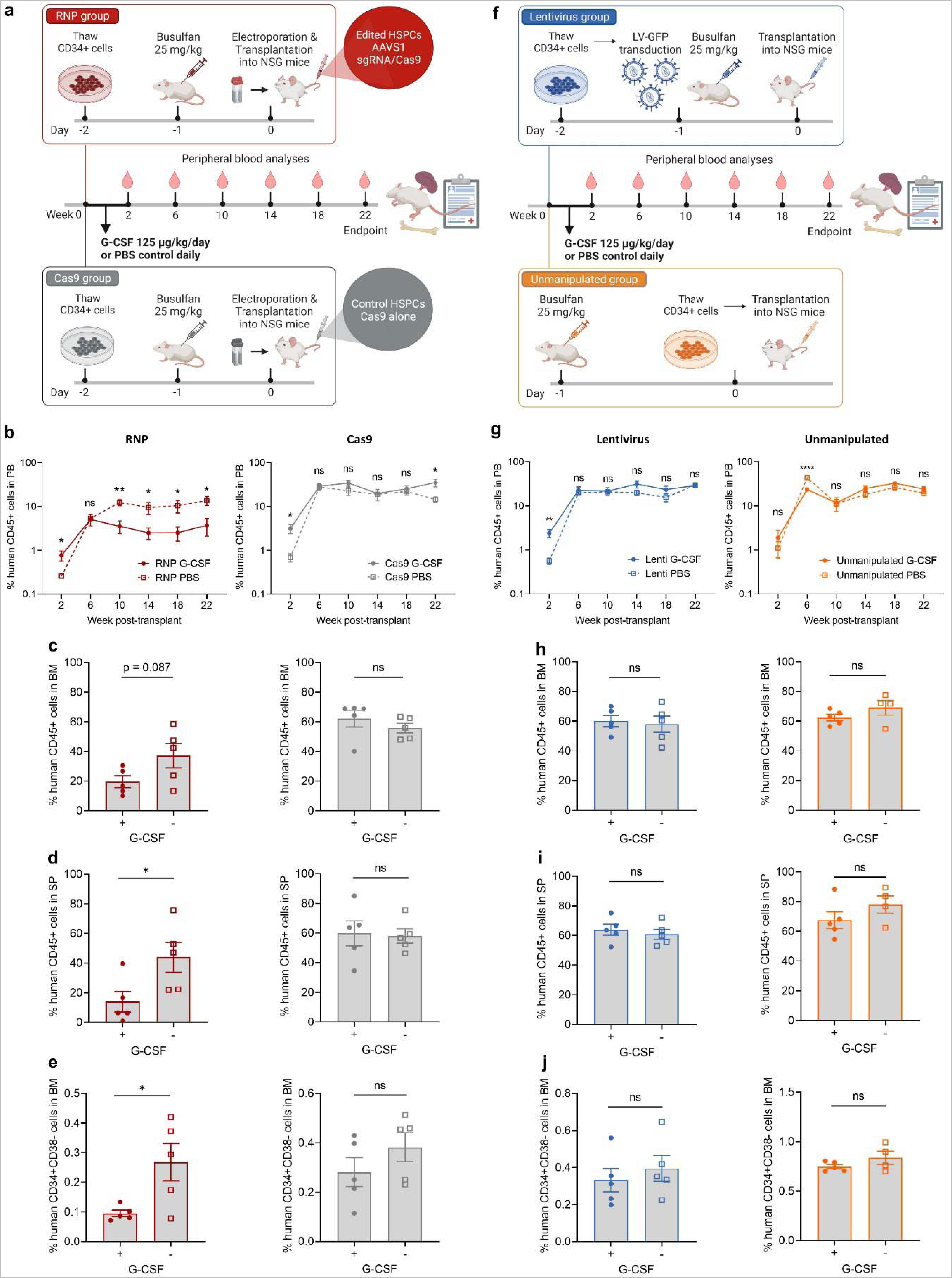
Post-transplant administration of G-CSF impedes long-term engraftment of CRISPR-Cas9 gene edited HSPCs but has minimal impact on lentiviral vector transduced or unmanipulated HSPCs. **a**, Experimental scheme: A total of 1 x 10^6^ human mobilized peripheral blood (MPB) CD34+ cells were electroporated with AAVS1-specific sgRNA/Cas9 ribonucleoprotein complexes (RNP group) or Cas9 alone (Cas9 group) and subsequently transplanted into NSG mice conditioned with busulfan. G-CSF or PBS was subcutaneously injected from post-transplant day 1 to 14 and hematopoietic reconstitution was compared between both groups (n = 5 mice/group). **b**, Percentages of human CD45+ cells in the peripheral blood (PB). **c**, Percentages of human CD45+ cells in the bone marrow (BM). **d**, Percentages of human CD45+ cells in the spleen (SP). **e**, Percentages of LT-HSC enriched populations (CD34+CD38-) in the BM. **f**, Experimental scheme: A total of 1 x 10^6^ human MPB CD34+ cells transduced with lentiviral vectors expressing GFP (lentivirus group) or unmanipulated human MPB CD34+ cells (unmanipulated group) were transplanted into busulfan conditioned NSG mice. G-CSF or PBS was subcutaneously injected from post- transplant day 1 to 14 and hematopoietic reconstitution was compared between both groups (n = 4-5 mice/group). **g**, Percentages of human CD45+ cells in the PB. **h**, Percentages of human CD45+ cells in the BM. **i**, Percentages of human CD45+ cells in the SP. **j**, Percentages of HSC enriched populations (CD34+CD38-) in the BM. In all panels, data are displayed as mean ± standard error of the mean (SEM), and two-sided unpaired t-test was used. ns, not significant, * p ≤ 0.05, ** p ≤ 0.01, **** p ≤ 0.0001.

Consistent with the clinical observation that G-CSF shortens time to neutrophil recovery after autologous HSPC transplantation in human subjects, G-CSF initially increased human CD45+ cells in the peripheral blood (PB) at 2 weeks post-transplant (Fig. 1b) by enhancing human CD13+ myeloid cell proliferation/differentiation from committed progenitors in both RNP and Cas9 groups (Extended Data Fig. 1b). However, starting at 10 weeks post-transplant when hematopoiesis begins to emerge from the most primitive HSPC populations, administration of G-CSF resulted in a 3-to 4-fold reduction in PB human cell engraftment compared to untreated mice in the RNP group, whereas no statistically significant differences were observed for this parameter between G-CSF treated and untreated mice in the Cas9 group (Fig. 1b). Similarly, BM and splenic engraftment in the G-CSF treated animals was 2- and 3-fold lower, respectively, at the 22-week endpoint analysis in the RNP group (Fig. 1c,d). The editing efficiency and lineage composition within human CD45+ cells were globally comparable between G-CSF treated and untreated mice in both RNP and Cas9 groups (Extended Data Fig. 1a-c). Importantly, percentages of LT-HSC enriched CD34+CD38- cell populations were 2.8-fold lower within the BM of G-CSF treated mice relative to untreated animals in the RNP group (Fig. 1e). In contrast, BM and splenic engraftment as well as percentages of BM CD34+CD38- cells were all comparable between G-CSF treated and untreated mice in the Cas9 group (Fig. 1c-e). Together, these data indicate that administration of G-CSF post-transplantation impedes engraftment of CRISPR-Cas9 gene edited HSPCs and this effect is attributable to cellular processes specifically triggered by gene editing rather than global toxicities from electroporation or Cas9 delivery.

### Post-transplant G-CSF administration has minimal impact on long-term engraftment of lentiviral vector transduced or unmanipulated HSPCs

We next investigated the functional consequences of post-transplant G-CSF administration on human CD34+ cells genetically engineered by transduction with lentiviral vectors (LV) and on control unmanipulated HSPCs (Fig. 1f). In the first group (designated “lentivirus group”), human MPB CD34+ cells were pre-stimulated overnight and subsequently transduced for 24 hours with LV expressing GFP at a multiplicity of infection (MOI) of 25. We then transplanted each busulfan-conditioned NSG mouse with the outgrowth of 1 x 10^6^ CD34+ cells seeded at the start of culture. In the second group (designated “unmanipulated group”), we similarly transplanted 1 x 10^6^ fresh human MPB CD34+ cells per NSG mouse conditioned with busulfan. Animals from both groups received daily subcutaneous injections of G-CSF or PBS during the first two weeks following transplantation, and hematopoietic reconstitution was compared between G-CSF treated and untreated animals.

Unlike our previous findings using CRISPR-Cas9 gene edited cells, we observed a comparable long-term PB human cell engraftment between G-CSF treated and untreated animals in both lentivirus and unmanipulated groups (Fig. 1g). Likewise, there were no statistically significant differences in levels of BM and splenic human CD45+ cell chimerism or percentages of LT-HSC enriched CD34+CD38- cells within the BM between G-CSF treated and untreated mice at the endpoint analysis (Fig. 1h-j). To exclude the possibility that transplantation of saturating numbers of human MPB CD34+ cells into NSG mice (i.e., 1 x 10^6^ cells/mouse) could have mitigated the G-CSF effects on engraftment, we included an additional “unmanipulated low-cell-dose group” (i.e., 1 x 10^5^ cells/mouse). Consistent with the high-cell-dose data, G-CSF had no appreciable impact on human CD45+ cell engraftment in the PB, BM and spleen at 22 weeks post-transplantation (Extended Data Fig. 2a-d). The lineage composition within human CD45+ cells was also comparable between G-CSF treated and untreated mice in all groups (Extended Data Fig. 2e). In the lentivirus group, *in vivo* GFP percentages within the CD13+ myeloid, CD20+ B lymphoid, CD3+ T lymphoid and CD34+ progenitor cell compartments were similar in the PB, BM and spleen of mice treated or not with G-CSF (Extended Data Fig. 2f). Collectively, these findings indicate that post-transplant administration of G-CSF has minimal impact on long-term engraftment of LV transduced or unmanipulated HSPCs.

### Post-transplant administration of G-CSF reduces frequencies and absolute numbers of CRISPR-Cas9 gene edited HSPCs

We next sought to quantitatively measure the impact of post-transplant G-CSF administration on CRISPR-Cas9 edited LT-HSCs using a limiting-dilution secondary transplantation assay. Human CD45+ cells isolated from the BM of primary mice in both RNP and Cas9 groups were serially transplanted into secondary immunodeficient NBSGW mice at limiting dilution and BM engraftment was analyzed at 20 weeks after transplantation (total period of engraftment: 42 weeks) (Fig. 2a). In the Cas9 group, similar percentages of mice transplanted with human CD45+ cells derived from primary animals treated with or without G-CSF met secondary multilineage BM engraftment criteria (47% vs. 57%, respectively), whereas administration of G-CSF to primary mice substantially reduced secondary engraftment in the RNP group (21% vs. 53%, respectively) (Fig. 2b). Extreme limiting dilution analysis (ELDA) indicated a 5.3-fold lower frequency of LT-HSCs within human CD45+ cells in the G-CSF treated RNP group relative to the untreated RNP group (p = 0.009) (Fig. 2c,d). In contrast, frequencies of LT-HSCs were comparable between G-CSF treated and untreated animals within the Cas9 group (p = 0.538) (Fig. 2c,d). To account for differences in human BM chimerism between primary treatment groups (Fig. 1c), we also quantified the absolute numbers of LT-HSCs within human CD45+ cells harvested from 4 hindlimb and 2 pelvic bones of transplanted animals. Although G-CSF treatment after primary transplantation only minimally reduced (1.1-fold) LT-HSC numbers compared to untreated mice in the Cas9 control group, we observed a substantial net decrease (11.1-fold) in the absolute numbers of LT-HSCs when primary mice from the RNP group were treated with G-CSF (Fig. 2d). Collectively, these data indicate that gene edited LT-HSCs are quantitatively diminished by post-transplant administration of G-CSF.

**Fig. 2.**
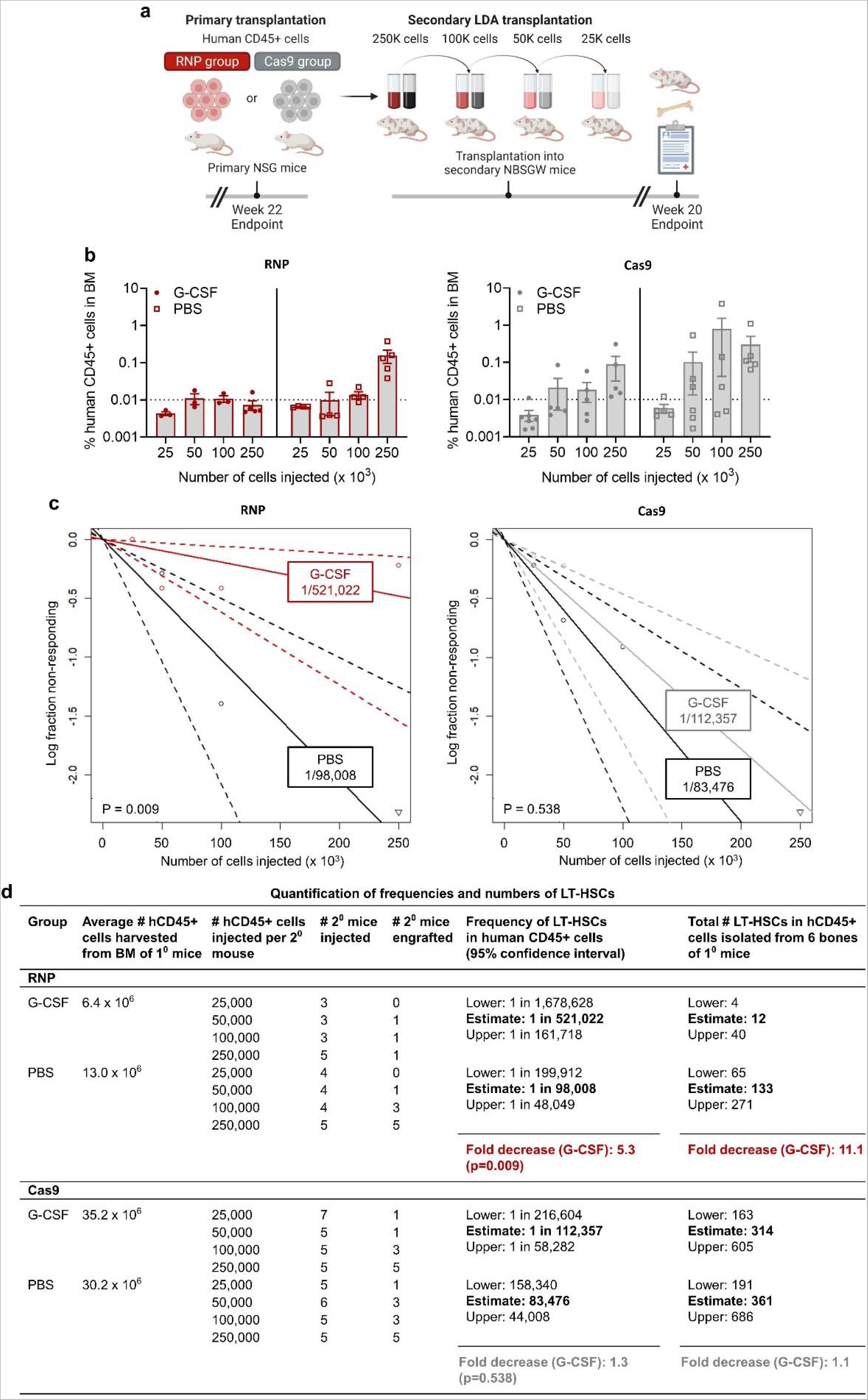
Post-transplant administration of G-CSF reduces frequencies and absolute numbers of gene edited LT-HSCs. **a,** Experimental scheme: Human CD45+ cells obtained from the bone marrow (BM) of primary mice in both RNP and Cas9 groups were serially transplanted at limiting dilution into secondary immunodeficient NBSGW mouse following a low dose busulfan conditioning. Human CD45+ cell engraftment within the BM was analyzed at 20 weeks post-transplant. **b**, Percentages of human CD45+ cells in the BM after secondary transplantation across multiple cell input doses (n = 3-7 mice/cell dose). Dashed line indicates cutoff for percentages of human CD45+ cells (>0.01%), including both myeloid (CD13+) and lymphoid lineages (CD20+), considered positive for calling secondary engraftment. **c**, Semilogarithmic plots of self-renewing LT-HSC frequencies within human CD45+ cells after secondary transplantation for the RNP and Cas9 groups. Solid lines indicate the best-fit linear model for each dataset. Dotted lines represent 95% confidence intervals. **d**, Quantification of LT-HSC frequencies and absolute numbers using a limiting-dilution secondary transplantation assay. Numbers of LT-HSCs were computed by arithmetic multiplication of the LT-HSC frequencies (column 6) by the average pooled numbers of hCD45+ cells harvested from the hindlimb bones of 5 primary mice transplanted with an equal number of input CD34+ cells (1 x10^6^ /mouse) for each group (column 2). P-values were calculated by extreme limiting dilution statistics. In panels b, data are displayed as mean ± standard error of the mean (SEM).39 | Page

### G-CSF exacerbates the p53 DDR activated by Cas9-induced DNA DSBs and transient p53 inhibition mitigates the G-CSF effects in gene edited human HSPCs

To gain insights into the potential mechanisms underlying the G-CSF-mediated engraftment defect of gene edited HSPCs, we first confirmed that surface G-CSFR is highly expressed on human MPB CD34+ cells, including a subset enriched in LT-HSCs (CD34+CD38-CD90+CD45RA-CD49f+). Additionally, HSPCs bearing the G-CSF receptor can sustain long-term multilineage hematopoietic reconstitution in serial xenotransplantation assays, suggesting a potential direct effect of G-CSF on these cells (Extended Data Fig. 3a-f).

We next queried whether G-CSF might influence the DDR mediated by the regulatory protein p53, the predominant cellular response activated by nuclease-induced DNA DSBs in human HSPCs^10–12, 31^. We cultured Cas9- and RNP-electroporated human CD34+ cells with and without G-CSF and quantified the expression level of the p53 DDR effector protein p21 by RT-qPCR analysis (Fig. 3a). Consistent with previous studies^10–12, 31^, we observed an upregulation of p21 gene expression in the RNP group relative to the Cas9 control group in the absence of G-CSF (Fig. 3b). Notably, p21 gene expression was further upregulated by addition of G-CSF during culture in the RNP group but unaltered in the Cas9 control group (Fig. 3b). To substantiate this observation, we transiently inhibited the p53 pathway by co-electroporating GSE56 mRNA during the gene editing procedure (designated “RNP/GSE56 group”). GSE56 is an mRNA encoding for a dominant negative p53 truncated polypeptide^28^ and co-electroporation of GSE56 was previously shown to provide a transient p53 inhibition (∼ 24 hours post-editing) and safely increase the yield of clonogenic and repopulating gene edited HSPCs^10–12^. A significantly lower level of p21 gene expression was observed in the RNP/GSE56 group relative to the RNP group, validating the co-electroporation of GSE56 mRNA as an effective strategy to prevent activation of the p53 pathway during gene editing in our experimental setting (Fig. 3b). Remarkably, addition of GSE56 completely offset the upregulation of p21 gene expression mediated by G-CSF in gene edited CD34+ HSPCs (Fig. 3b).

**Fig. 3.**
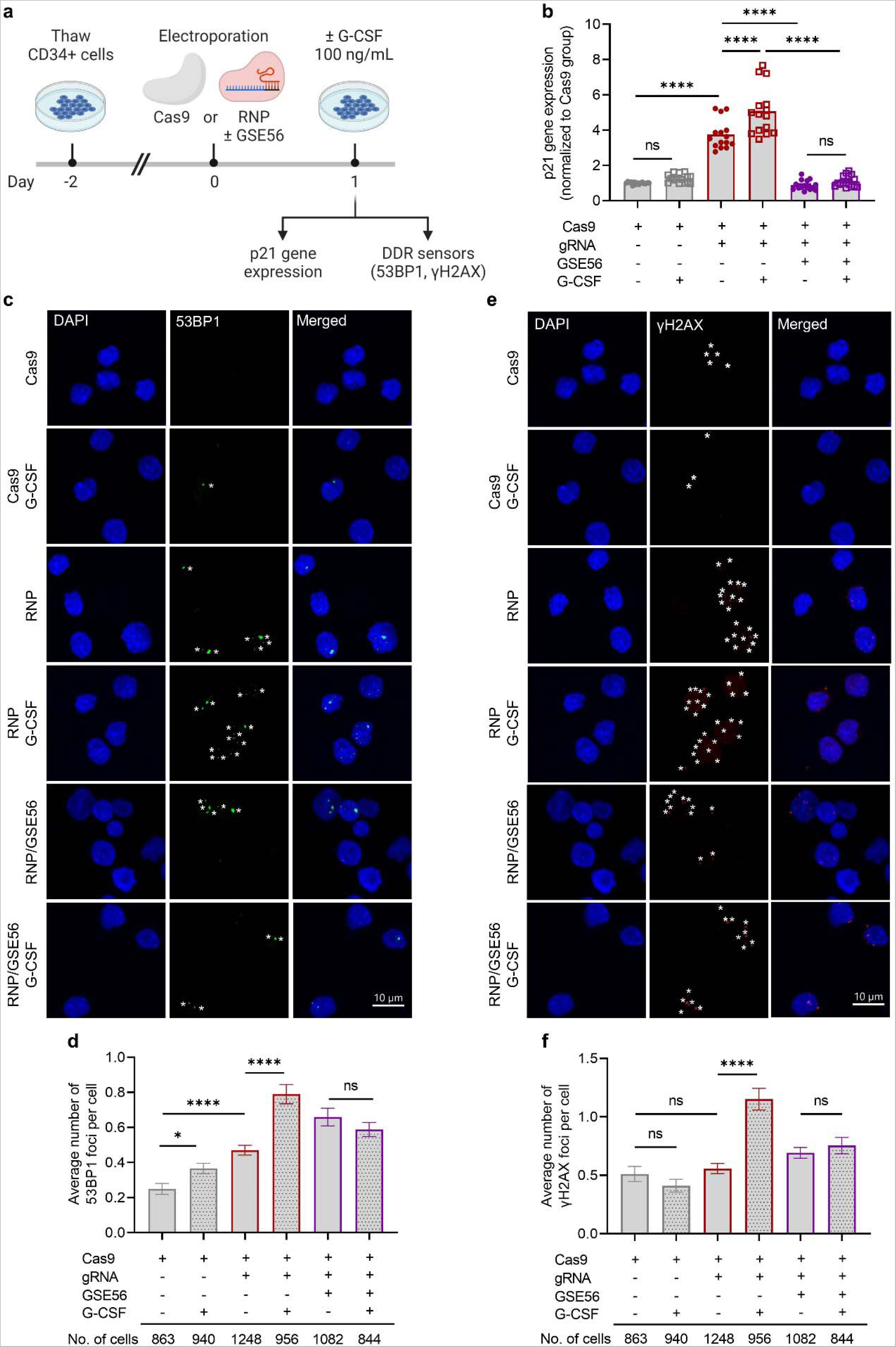
G-CSF exacerbates the p53 DDR activated by Cas9-induced DNA DSBs and transient p53 inhibition mitigates the G-CSF effects in human HSPCs. **a,** Experimental scheme. Pre-cultured human MPB CD34+ cells were electroporated with Cas9 alone (Cas9 group), AAVS1-specific sgRNA/Cas9 RNP (RNP group), or RNP in the presence of GSE56 mRNA (RNP/GSE56 group). Edited CD34+ cells were further cultured for 4 hours in the presence or absence of G-CSF at a dose of 100 ng/mL. Expression of p21 and DNA damage response (DDR) sensors were measured by RT-qPCR and confocal microscopy, respectively, at 28 hours post-electroporation. **b**, Relative p21 gene expression, as measured by RT-qPCR (n = 15 technical replicates/group, from 5 independent donors). **c**, Representative confocal images of DAPI (blue) and DNA damage response sensor 53BP1 (green) within each group of human CD34+ HSPCs. Asterisks indicate 53BP1 foci. Scale bar represents 10 µm. **d**, Average numbers of 53BP1 foci per cell (n = 844-1248 cells/group, from 3 independent donors). **e**, Representative confocal images of DAPI (blue) and DNA damage response sensor γH2AX (red) within each group of human CD34+ HSPCs. Asterisks indicate γH2AX foci. Scale bar represents 10 µm. **f**, Average numbers of γH2AX foci per cell (n = 844-1248 cells/group, from 3 independent donors). In panel **b**, data are displayed as mean ± standard error of the mean (SEM) and one way ANOVA with Tukey’s multiple comparison test was used. In panels **d** and **f**, data are displayed as mean ± SEM and Kruskal-Wallis test with Dunn’s multiple comparison test was used. ns, not significant, * p ≤ 0.05, **** p ≤ 0.0001.

To corroborate these findings, we tracked DDR factors recruited at sites of DNA DSBs by quantifying microscopically visible 53BP1 and γH2AX subnuclear foci in Cas9-, RNP- and RNP/GSE56-electroporated human CD34+ cells cultured with and without G-CSF (Fig. 3a). In the absence of G-CSF, we observed a significant increase in the number of 53BP1 foci per cell in the RNP group relative to the Cas9 group, consistent with the known p53-mediated DDR induced by nuclease activity (Fig. 3c,d). However, the number of γH2AX foci per cell was similar between the RNP and Cas9 groups, in keeping with a previous study indicating that γH2AX signals can be induced by *ex vivo* culture itself and may be a less sensitive marker for detection of nuclease activity than 53BP1 signals^10^ (Fig. 3e,f). Nevertheless, supplementing the culture medium with G-CSF significantly increased the number of both 53BP1 and γH2AX foci per nucleus relative to untreated cells in the RNP group, whereas G-CSF had minimal or no impact in Cas9- and RNP/GSE56-treated cells (Fig. 3c-f). We observed a similar trend in response to G-CSF when percentages of 53BP1 and γH2AX foci-bearing cells were measured in each group, with most cells displaying one or two foci per nucleus independent of treatment (Extended Data Fig. 5a,b).

Taken together, these data indicate that G-CSF exacerbates the p53 DNA damage response triggered by Cas9-induced DNA DSBs via G-CSFR-linked cell intrinsic mechanisms, and transient inhibition of the p53 pathway mitigates this effect in gene edited HSPCs.

### G-CSF transcriptionally modulates the p53 and cell cycle checkpoint pathways in gene edited human HSPCs with a more prominent impact in LT-HSCs

To understand the deleterious impact of G-CSF at single cell level and across HSPC subpopulations, we performed Cellular Indexing of Transcriptomes and Epitopes sequencing (CITE-seq) of Cas9-, RNP- and RNP/GSE56-electroporated human CD34+ cells treated with G-CSF or PBS after their transplantation into NSG mice (Fig. 4a). CITE-seq was implemented with a combination of antibodies customarily accepted to define a CD34+CD38-CD90+CD45RA- CD49f+ cell population highly enriched in LT-HSCs^32, 33^. Additionally, to eliminate a batch effect during analysis, all experimental samples were pooled and sequenced concurrently by applying anti-human hashtag antibodies before CITE-seq library preparation. We used Seurat R package to perform quality control and demultiplex the pooled sequenced data into individual experimental groups for comparative analyses. Summary statistics are shown in Extended Data Fig. 5a-d.

**Fig. 4.**
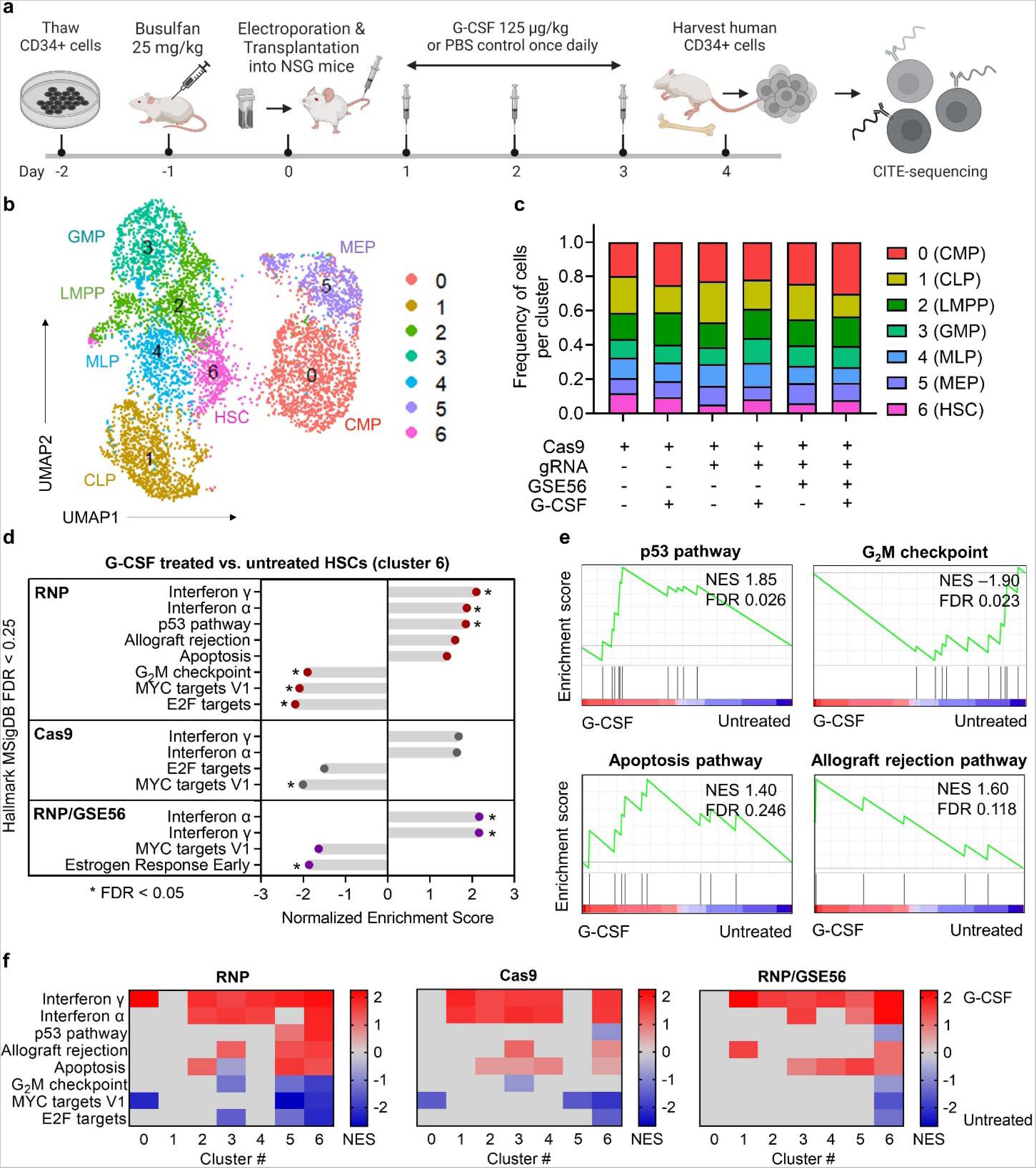
G-CSF transcriptionally modulates the p53 and cell cycle checkpoint pathways in gene edited human HSPCs with a more prominent impact in LT-HSCs. **a**, Experimental scheme for cell preparation and CITE-seq. Human MPB CD34+ cells treated with Cas9 alone (Cas9 group), AAVS1-specific sgRNA/Cas9 RNP (RNP group), or RNP in the presence of GSE56 mRNA (RNP/GSE56 group) were transplanted into NSG mice after busulfan conditioning. G-CSF or PBS was subcutaneously injected from post-transplant day 1 to 3. Human CD34+ cells were isolated from the murine bone marrow at the end of the treatment and processed for CITE-seq analysis. **b**, Merged UMAP visualization of CITE-seq data for CD34+ cells from each experimental group. A total of seven cell clusters were identified and annotated as common myeloid progenitors (CMP, cluster 0), common lymphoid progenitors (CLP, cluster 1), lymphoid-primed multipotent progenitors (LMPP, cluster 2), granulocyte-monocyte progenitors (GMP, cluster 3), multilymphoid progenitors (MLP, cluster 4), megakaryocytic- erythroid progenitors (MEP, cluster 5) and hematopoietic stem cell (HSC, cluster 6). **c**, Stacked bar plots showing frequency of the identified clusters across all samples. **d**, Gene set enrichment analysis (GSEA) using ranked differentially expressed gene sets obtained by comparing G-CSF treated and untreated HSC clusters against the Hallmark gene set of the Molecular Signature database (MSigDB). Pathways with statistically significant differences (i.e., false discovery rate (FDR) < 0.25) are shown. **e**, Enrichment plots illustrating the relative expression of genes associated with p53, allograft rejection, apoptosis and G2M checkpoint pathways. NES, normalized enrichment score; FDR, false discovery rate. **f**, Heatmap displaying NES of pathways with statistically significant differences in HSC cluster 6 of the RNP group across all clusters (0-6).

We clustered and visualized transcriptome sequencing data of single CD34+ cells from each experimental group in two-dimensional uniform manifold approximation and projection (UMAP). We identified seven distinct clusters (clusters 0 to 6) (Fig. 4b), with a similar distribution of cells per cluster between all experimental groups (Fig. 4c). We assigned each CD34+ cell cluster to a distinct HSPC subpopulation, including common myeloid progenitors (CMP, cluster 0), common lymphoid progenitors (CLP, cluster 1), lymphoid-primed multipotent progenitors (LMPP, cluster 2), granulocyte-monocyte progenitors (GMP, cluster 3), multilymphoid progenitors (MLP, cluster 4), megakaryocytic-erythroid progenitors (MEP, cluster 5) and a single LT-HSC population (cluster 6). CITE-seq cluster assignment was computed by associating cluster-specific transcripts with established lineage markers (Extended Data Fig. 5e) and HSC transcriptomic signatures (Extended Data Fig. 5f), and subsequently confirmed by pseudotime ordering analysis (Extended Data Fig. 5g) and patterns of surface expression for CD38, CD45RA, CD90 and CD49f markers in each cluster (Extended Data Fig. 5h).

We first assessed the relative impact of G-CSF on the LT-HSC population by gene set enrichment analysis (GSEA) using ranked differentially expressed gene sets obtained by comparing G-CSF treated and untreated cells within LT-HSC cluster 6. We observed several differentially expressed genes between experimental conditions (Supplementary Table 1). GSEA based on the Hallmark gene set of the Molecular Signature database revealed an upregulation of interferon α/γ inflammatory responses and a downregulation of E2F and MYC targets in LT-HSCs exposed to G-CSF relative to control LT-HSCs in all groups (i.e., RNP, Cas9 and RNP/GSE56), suggesting that administration of G-CSF alone modulated these pathways in human LT-HSCs *in vivo* independent of the gene editing process. In contrast, four pathways were modulated by G-CSF only in RNP-treated LT-HSCs, including the p53, G_2_M checkpoint, apoptosis and allograft rejection (i.e., immune rejection/defense) pathways (Fig. 4d,e). In agreement with our RT-qPCR analysis (Fig. 3b), heightened upregulation of the p53 pathway was the predominant response to G-CSF in gene-edited LT-HSCs. The p53 DDR encompasses signal transduction pathways that effect cell cycle checkpoint arrest to prevent entry into mitosis until DNA damage is repaired and/or cellular apoptosis if genomic integrity is irreparably compromised. The G_2_M DNA damage checkpoint was downregulated in gene edited LT-HSCs exposed to G-CSF, suggesting that damaged cells may have arrested in an earlier cell cycle phase for repair or death, or may be allowed to proliferate unrepaired under the stimulatory influence of G-CSF. In support of the first possibility, the apoptosis pathway was exclusively upregulated in the RNP group by G-CSF treatment although the increase was marginal and did not reach statistical significance. Additionally, consistent with the emerging concept of a crosstalk between DDR and the immune system^34^, the allograft rejection pathway was also stimulated by G-CSF in the RNP group only but, as for apoptosis, the extent of stimulation was limited. (Fig. 4d,e). Notably, when the top differentially expressed gene sets in LT-HSC cluster 6 were examined across all cell clusters in both RNP and Cas9 groups, we observed a more prominent G-CSF-mediated modulation of these pathways within transcriptionally defined LT-HSCs compared to lineage-restricted progenitor cell populations in clusters 0–5 (Fig. 4f, Supplementary Table 2).

To substantiate the observed transcriptional impact of G-CSF on p53-mediated DDR and associated pathways in gene edited human HSPCs, we compared differentially expressed gene sets in G-CSF treated and untreated CD34+ cells subjected to transient p53 inhibition with GSE56 during the gene editing procedure. Transient attenuation of the p53 pathway mitigated the G-CSF-dependent transcriptional response in p53, cell cycle checkpoint, apoptosis and allograft rejection gene sets within LT-HSC cluster 6 and, to a lesser degree, some of the progenitor populations in clusters 0-5 (Fig. 4d,f, Supplementary Table 1).

Taken together, these data indicate that G-CSF treatment of gene edited human HSPCs primarily modulates transcriptional activity of the p53 DNA damage response and cell cycle checkpoint pathways, and modestly induces gene sets involved in apoptosis and immune rejection/defense. Importantly, G-CSF has a more pronounced impact on the transcriptionally defined LT-HSC cellular subset relative to committed progenitor cell clusters.

### G-CSF promotes S-phase cell cycle arrest in gene edited LT-HSCs and has a limited impact on cell survival

To independently confirm the p53-associated effects of G-CSF on cell cycle progression and apoptosis of gene edited HSPCs, we assayed by BrdU incorporation and AnnexinV staining, respectively, populations of human CD34+CD38- cells (enriched in LT-HSCs) and CD34+CD38+ cells (hematopoietic progenitors) isolated from the BM of xenotransplanted mice treated with G-CSF or control PBS (Fig. 5a). In hematopoietic progenitors, the cell cycle status was globally similar between G-CSF treated and untreated cells in all groups (Fig. 5b,c). Notably, G-CSF treatment *in vivo* increased the frequency of cells in S-phase and decreased the frequency of cells in G_2_M-phase within the LT-HSC enriched CD34+CD38- cell fraction in the RNP group (Fig. 5d,e). In contrast, the proportion of cells in S- and G_2_M-phases within CD34+CD38- hematopoietic populations remained unchanged after G-CSF treatment in the Cas9 and RNP/GSE56 groups (Fig. 5d). Lastly, consistent with the limited upregulation of apoptosis pathways delineated by CITE-seq analysis (Fig. 4d), a positive trend in Annexin V+ cells was also observed exclusively within RNP-treated CD34+CD38- populations exposed to G-CSF relative to PBS controls; the rate of apoptosis was otherwise unaltered by G-CSF in both progenitors and LT-HSCs within Cas9, RNP and RNP/GSE56 groups (Fig. 5f,g).

**Fig. 5.**
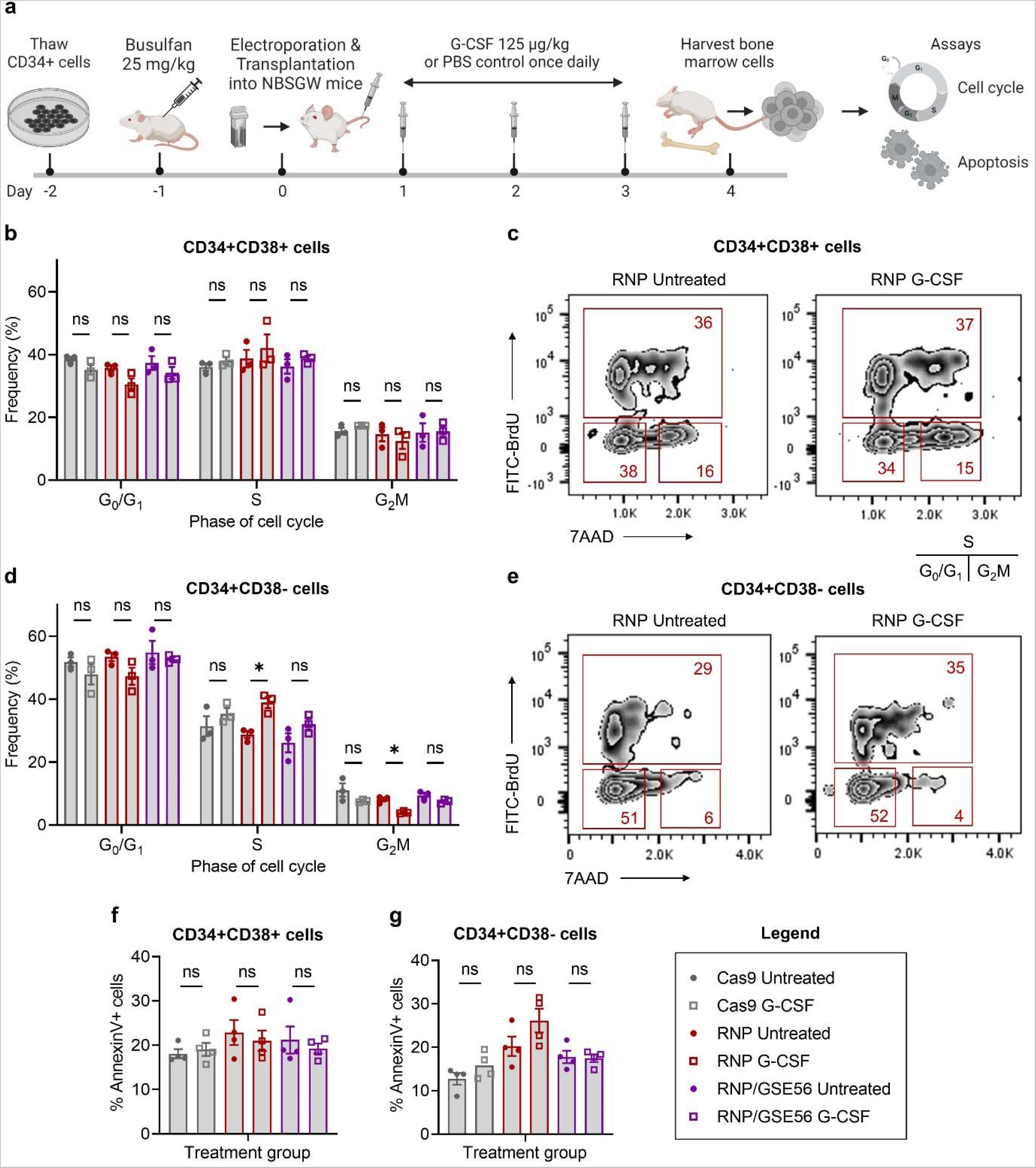
G-CSF promotes cell cycle arrest in gene edited LT-HSCs. **a,** Experimental scheme for cell cycle analysis and apoptosis assay. Human MPB CD34+ cells treated with Cas9 alone (Cas9 group), AAVS1-specific sgRNA/Cas9 RNP (RNP group), or RNP in the presence of GSE56 mRNA (RNP/GSE56 group) were transplanted into NBSGW mice after busulfan conditioning. G-CSF or PBS was subcutaneously injected from post-transplant day 1 to 3 and the murine bone marrow (BM) was harvested for flow cytometric analyses. **b**, Percentages of CD34+CD38+ hematopoietic populations in indicated cell cycle phases (n = 3 independent donors). **c**, Representative flow cytometry profiles of CD34+CD38+ hematopoietic populations in the RNP group with or without G-CSF treatment. **d**, Percentages of CD34+CD38- hematopoietic populations in indicated cell cycle phases (n = 3 independent donors). **e**, Representative flow cytometry profiles of CD34+CD38- hematopoietic populations in the RNP group with or without G-CSF treatment. **f-g**, Percentages of AnnexinV+ cells within CD34+CD38+ (f) and CD34+CD38- (g) hematopoietic populations (n = 4 independent donors). In panels b, d, f and g, data are presented as mean ± standard error of the mean (SEM). Two- sided unpaired t-test with Bonferroni correction was used. ns, not significant, * p ≤ 0.05.

From these data, we infer that G-CSF-mediated DDR induces S-phase arrest with a modest impact on cellular survival and transient p53 inhibition attenuates this process in gene edited LT-HSCs.

### Transient p53 inhibition during gene editing mitigates the negative impact of G-CSF on gene edited LT-HSC function

We next used xenotransplantation assays to investigate whether attenuation of p53- mediated DDR with GSE56 could alter engraftment of gene edited LT-HSCs *in vivo*. Human MPB CD34+ cells isolated from the same healthy donor used in the previous studies were pre-stimulated for 2 days and subsequently electroporated with AAVS1-specific sgRNA/Cas9 RNP complexes in the presence of GSE56. The outgrowth of 1 x 10^6^ CD34+ cells seeded at the start of culture was transplanted into each busulfan-conditioned NSG mouse, and the animals received G-CSF or PBS subcutaneously from post-transplant day 1 to 14 (Fig. 6a). In the RNP group, an editing efficiency of 68% in bulk CD34+ HSPCs was measured by T7 endonuclease I (T7EI) cleavage of target site PCR amplicons (Extended Data Fig. 6a).

**Fig. 6.**
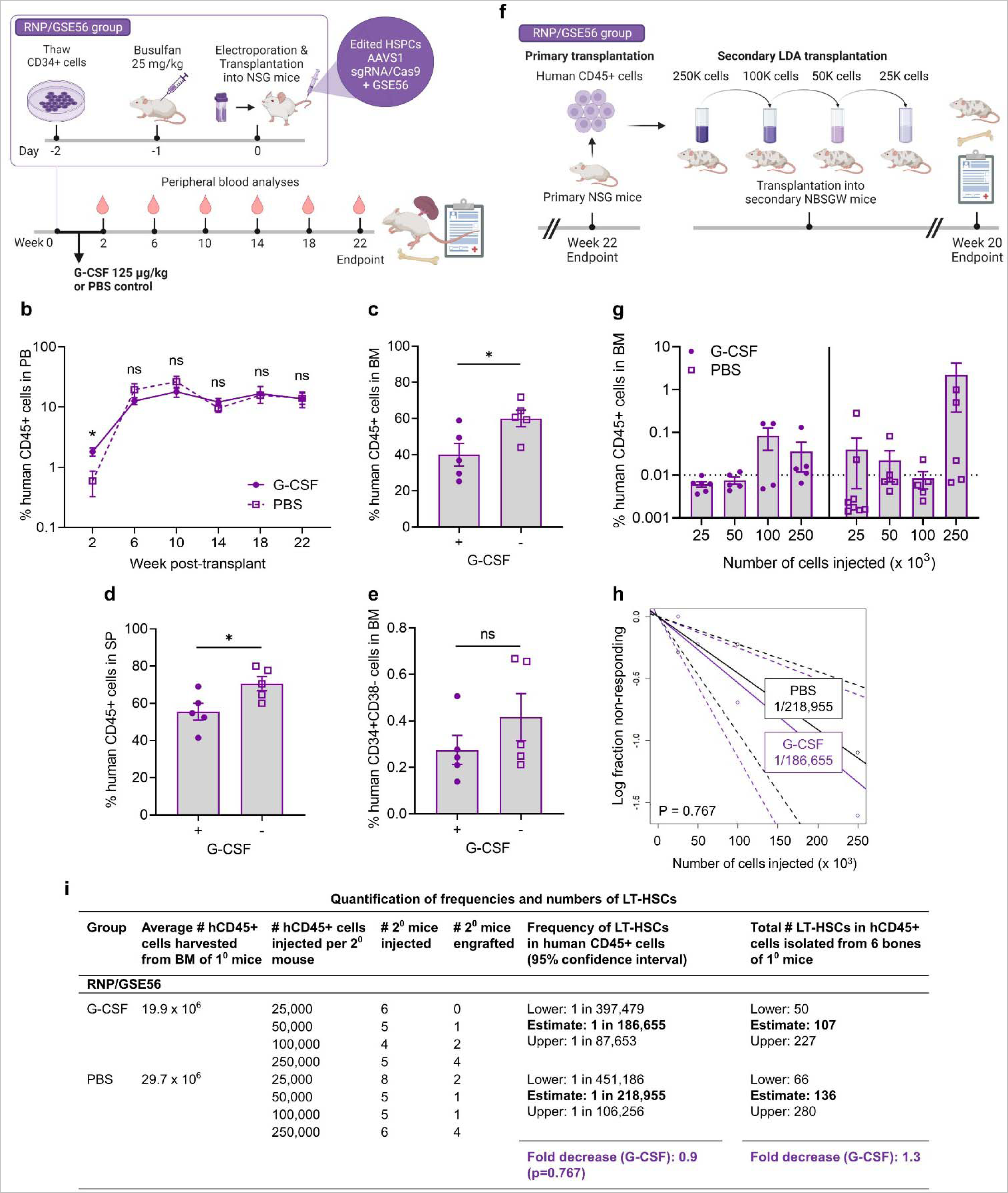
Transient p53 inhibition during gene editing mitigates the negative impact of G- CSF on gene edited LT-HSC function. **a**, Experimental scheme: Human MPB CD34+ cells were electroporated with AAVS1-specific sgRNA/Cas9 ribonucleoprotein (RNP) complexes in the presence of GSE56 mRNA and subsequently transplanted into NSG mice after busulfan conditioning. G-CSF or PBS was subcutaneously injected from post-transplant day 1 to 14 and hematopoietic reconstitution was compared between both groups (n = 5 mice/group). **b**, Percentages of human CD45+ cells in the peripheral blood (PB) of primary mice. **c**, Percentages of human CD45+ cells in the bone marrow (BM) of primary mice. **d,** Percentages of human CD45+ cells in the bone marrow (BM) of primary mice. **e**, Percentages of LT-HSC enriched populations (CD34+CD38-) in the BM of primary mice. **f**, Experimental scheme: Human CD45+ cells obtained from the BM of primary mice in the RNP/GSE56 group were serially transplanted into immunodeficient NBSGW mice at limiting dilution following a low dose busulfan conditioning and BM engraftment was analyzed at 20 weeks post-transplant. **g**, Percentages of human CD45+ cells in the BM after secondary transplantation across multiple cell input doses (n = 4-8 mice/cell dose). Dashed line indicates cutoff for percentages of human CD45+ cells (>0.01%), including both myeloid (CD13+) and lymphoid lineages (CD20+), considered positive for calling secondary engraftment. **h**, Semilogarithmic plots of self-renewing LT-HSC frequencies within human CD45+ cells after secondary transplantation for the RNP/GSE56 group. Solid lines indicate the best-fit linear model for each dataset. Dotted lines represent 95% confidence intervals. **i.** Quantification of LT-HSC frequencies and absolute numbers using a limiting-dilution secondary transplantation assay. Numbers of LT-HSCs were computed by arithmetic multiplication of the LT-HSC frequencies (column 6) by the average pooled numbers of hCD45+ cells harvested from the hindlimb bones of 5 primary mice transplanted with an equal number of input CD34+ cells (1 x 10^6^ /mouse) for each group (column 2). P-values were calculated by extreme limiting dilution statistics. In panels **b-e** and **g**, data are displayed as mean ± standard error of the mean (SEM) and two-sided unpaired t-test was used. ns, not significant, * p ≤ 0.05.

In agreement with our previous observations, G-CSF treatment augmented human CD45+ cell chimerism at 2 weeks post-transplant (Fig. 6b) by enhancing human CD13+ myeloid cell proliferation/differentiation (Extended Data Fig. 6b). Interestingly, in contrast to our previous findings with RNP-edited cells (Fig. 1b), we observed a comparable long-term PB human cell engraftment between G-CSF treated and untreated mice in the RNP/GSE56 group, suggesting that transient p53 inhibition attenuated the negative effects of G-CSF on gene edited repopulating HSPCs upon primary transplantation (Fig. 6b). Likewise, the magnitude of differences in BM and splenic engraftment levels between G-CSF treated and untreated mice was ameliorated in the RNP/GSE56 group (Fig. 6c,d) compared to studies conducted in the absence of GSE56 (Fig. 1c,d). The editing efficiency and lineage composition within human CD45+ cells were globally comparable between G-CSF treated and untreated mice (Extended Data Fig. 6a-c). Importantly, the reduction in LT-HSC enriched CD34+CD38- cells within the BM of G-CSF treated vs. untreated mice previously observed in the RNP group (Fig. 1e) was alleviated by addition of GSE56 during the gene editing process (Fig. 6e).

To quantitatively assess the impact of transient p53 inhibition on gene edited LT-HSCs exposed to G-CSF, we performed a limiting-dilution secondary transplantation analysis of the RNP/GSE56 group (Fig. 6f). Although administration of G-CSF to primary mice substantially reduced secondary engraftment of gene edited HSPCs (Fig. 2b), this effect was reversed by addition of GSE56 during the gene editing procedure, as shown by similar percentages of mice meeting secondary multilineage BM engraftment criteria after G-CSF (35%) or control PBS treatment (33%) of primary recipient mice (Fig. 6g). Correspondingly, the previously observed 5.3-fold reduction in LT-HSC frequency caused by G-CSF administration in the RNP group (Fig. 2c,d) was mitigated by adjunct GSE56 mRNA during gene editing (Fig. 6h,i). Likewise, absolute numbers of LT-HSCs within human CD45+ cells harvested from 4 hindlimb and 2 pelvic bones of transplanted animals within the RNP/GSE56 group were only minimally diminished (1.3-fold) by G-CSF treatment (Fig. 6i), representing a substantial improvement relative to the observed 11.1-fold reduction in the absolute numbers of LT-HSCs when primary mice from the RNP group were treated with G-CSF (Fig. 2d). Collectively, these data indicate that transient p53 inhibition during gene editing largely rescues the loss of gene edited LT-HSCs mediated by post-transplantation administration of G-CSF and confirm the regulatory protein p53 as a central point of convergence for pathways induced by nuclease-based gene editing and G-CSF treatment.

## Discussion

In this study, we provide functional and molecular evidence that administration of G-CSF after transplantation of CRISPR-Cas9 nuclease gene edited HSPCs exacerbates the p53 DDR triggered by the gene editing process. This response is more robust in the primitive LT-HSC cellular subset and significantly impairs long-term hematopoietic reconstitution after transplantation. Thus, the p53 pathway represents a key point of convergence for nuclease-induced DNA damage and G-CSF treatment.

The observed toxicity of G-CSF on gene edited human HSPCs may be ascribed to robust activation of the p53 DDR pathway in the unique cellular context of CRISPR-induced DNA double-stranded breaks. This is suggested, in part, by reversal of the deleterious effects of G-CSF on gene edited HSPCs by dampening p53 expression with GSE56 during the gene editing procedure. The importance of counteracting DDR activation was also highlighted in previous investigations co-electroporating GSE56 mRNA during gene editing of human CD34+ cells to transiently lower DDR burden and alleviate clonogenic and repopulation defects in gene edited HSPCs^10–12^. Additionally, although exposure to lentivirus^35^ and Cas9 alone^36^ can also activate the p53 pathway, these manipulations combined with G-CSF administration did not provide a synergy sufficient to impair long-term engraftment of human HPSCs in our study. This observation is consistent with a threshold mechanism by which expression levels of p53 and its targets must be sufficient to induce growth arrest or apoptotic transcriptional programs^37^. The synergistic activation of p53 observed in HSPCs exposed to G-CSF and CRISPR-Cas9 RNPs is analogous to the enhanced p53 stimulation previously reported when HSPC gene editing was performed in conjunction with transduction of AAV6 DNA donor template delivery vectors^10–12^. In the future, it will be important to ascertain the impact of post-transplant G-CSF on human HSPCs subjected to emerging gene correction technologies that do not rely on DSB formation (i.e., base editing^38–40^ and prime editing^40–43^) or are independent of the cellular DDR (i.e., recombinases^40, 44–47^ and RNA-guided transposon systems^40, 47^). Globally, our findings further expand the body of evidence delineating p53 as a central hub in a molecular network orchestrating a variety of DDR pathways, including cell cycle checkpoints, apoptosis and immune responses.

Further investigation is needed to fully unravel the mechanisms by which G-CSF intensifies the p53-mediated DDR in human HSPCs subjected to CRISPR-Cas9 gene editing and impairs their repopulating capacity. Key observations indicate that G-CSF acts, at least in part, in a cell-intrinsic manner to inhibit function of gene edited HSPCs. This hypothesis is primarily supported by the potent impact of G-CSF on p21 gene expression and accumulation of nuclear DDR foci in purified gene edited CD34+ HSPCs cultured without accessory cells *ex vivo*. The cell-intrinsic effect of G-CSF is likely mediated by activation of the G-CSF:G-CSFR signaling axis in HSPCs. Indeed, in agreement with a previous study^18^, we demonstrated robust expression of G-CSFR on human MPB CD34+ cells and long-term multilineage hematopoietic reconstitution after transplantation of HSPCs bearing the G-CSF receptor in primary and secondary xenotransplantation assays. Nevertheless, these findings do not exclude that some of the deleterious properties of G-CSF on HSPCs may be independent of G-CSFR stimulation. For instance, treatment with G-CSF was previously shown to induce a cell-autonomous HSPC repopulating defect through induction of Toll-like receptor (TLR) expression and signaling in murine models^48^. Although cell-intrinsic mechanisms likely influence how G-CSF negatively impacts nuclease-edited HSPCs, it is conceivable that microenvironmental and other cellular constituents also play a key role in this process. In particular, the stem cell mobilizing properties of G-CSF are well known in the clinic. This process does not depend on G-CSFR expression on HSPCs and is instead mechanistically linked to cell-extrinsic factors, including the loss of osteoblasts and reduced CXCL12 expression within the bone marrow niche^49–52^. Hence, an indirect mode of action via mobilization of gene edited HSPCs in recipient mice treated with G-CSF may also contribute to their loss of function *in vivo*.

Our results may also have considerable implications for gene therapies targeting hematopoietic stem and progenitor cells. Although G-CSF administration is standard of care after autologous HSPC transplantation^17^ and commonly used in the context of allogenic HSPC transplantation when benefits outweigh potential risks^53–57^, much less is known about its safety and efficacy after transplantation of genetically corrected autologous HSPCs. While the impact of G-CSF on gene edited HSPCs would optimally be addressed in clinical trials randomizing patients to receive G-CSF support or placebo during the post-transplant recovery phase, such trials are unlikely to be conducted due to ethical considerations and inherent biases. Here, using gold-standard surrogate LDA xenotransplantation assays, we observed a significant attrition of the LT-HSC pool recovered from mice treated with G-CSF after transplantation of gene edited human HSPCs. Because therapeutic success in gene therapy studies has been associated with high diversity of clonal reconstitution which correlates with the number of corrected HSPC clones infused back to the patient^58, 59^, administration of G-CSF post-transplantation may thus negatively impact clinical outcomes by reducing cell dose and clonal diversity in treated subjects. Although future studies will be necessary to precisely track individual clones long-term after transplantation, data presented in this study provide substantive evidence that G-CSF adjunct therapy should be used with caution in the context of autologous HSPC gene therapy clinical trials where DNA damaging programmable nucleases are utilized to manipulate human CD34+ HSPCs. In clinical settings such as leukocyte adhesion deficiency, chronic granulomatous disease or severe congenital neutropenia where G-CSF administration is frequently considered to facilitate recovery from severe neutropenia, transient p53 inhibition by addition of GSE56 mRNA at the time of editing may be considered to ensure improved clinical response. Ultimately, stakeholders need to carefully examine the risk/benefit balance of an intervention for individual patients, taking into consideration the clinical implications of prolonged neutropenia, reduced diversity of clonal reconstitution and overall toxicity of CRISPR-Cas9 gene editing approaches.

In sum, we provide new molecular and functional evidence suggesting the potential of G-CSF administration to exacerbate HSC toxicity triggered by nuclease-mediated DNA double stranded breaks. On the basis of these findings, we envision that the latent risks and benefits of G-CSF administration to individual patients will be carefully weighed as autologous HSPC gene editing studies are implemented in the clinic.

## Experimental Procedures

Reagents and resources used in this study are detailed in Supplementary Table 3.

### Isolation and culture of human CD34+ HSPCs

Human CD34+ HSPCs were obtained from a healthy male volunteer after informed consent in accordance with the Declaration of Helsinki, under an Institutional Review Board-approved clinical protocol (NCT00001529). Peripheral blood mobilization of HSPCs was induced via subcutaneous injection of 10 μg/kg G-CSF (Filgrastim, Amgen, Thousand Oaks, CA, USA) for 5 days followed by leukapheresis using a Cobe Spectra Apheresis System (Terumo BCT, Lakewood, CO, USA). The mononuclear cell concentrates were enriched in CD34+ HSPCs using a CliniMACS Plus instrument (Miltenyi Biotec, Gaithersburg, MD, USA) and cryopreserved prior to use. All human CD34+ HSPCs were cultured in StemSpan SFEM II medium (STEMCELL Technologies, Cambridge, MA, USA) supplemented with stem cell factor (SCF; 100 ng/mL), thrombopoietin (TPO; 100 ng/mL), Fms-like tyrosine kinase 3 ligand (Flt3- ligand; 100 ng/mL) and 1% penicillin-streptomycin (GIBCO, Grand Island, NY, USA), in the presence or absence of G-CSF at a dose of 100 ng/mL. Cells were cultured with 5% CO_2_ at 37°C.

### Electroporation of human HSPCs

Two days prior to electroporation, primary human HSPCs were thawed and plated at 0.5 x 10^6^ cells/mL in StemSpan SFEM II medium as detailed above. After 48 hours in culture, individual HSPC samples (∼3.5 x 10^6^ cells) were aliquoted into sterile microfuge tubes and pelleted at 3000 rpm for 5 min in a Sorvall Legend Micro 17 centrifuge (Thermo Fisher Scientific, Waltham, MA, USA). Cell pellets were resuspended in 100 μL MaxCyte electroporation buffer (MaxCyte, Inc., Gaithersburg, MD, USA) and electroporated (with or without Cas9 or Cas9 RNPs) in OC-100 x 3 Processing Assemblies (MaxCyte) using a MaxCyte GT electroporator and program HPSC34-3-OC. For gene editing, 20 μg of purified recombinant SpCas9 protein (PNA Bio, Inc., Thousand Oaks, CA, USA) was pre-complexed on ice for 20 min with 10 μg of the chemically modified sgRNA (bearing 2’-O methyl phosphorothioate-modified nucleotides at the first 3 and last 3 positions of the synthetic sgRNA) prior to addition to the sample. When transiently inhibiting the p53 pathway, GSE56 mRNA (NIH, NGRNA0210, Lot#200114) was added to SpCas9/sgRNA mixtures at a dose of 150 μg/mL. The cognate 20- mer incorporated into the individual sgRNAs to recognize the AAVS1 locus was as follows: 5’- GUUAAUGUGGCUCUGGUUCU-3’. “Cas9 only” samples received 20 ug SpCas9 in the absence of sgRNA. Electroporated cells were allowed to rest for 20 min at 37°C and then plated at a concentration of 0.5 x 10^6^ – 1.0 x 10^6^ cells/mL and incubated at 37°C in a 5% CO_2_ atmosphere or transplanted into NSG or NBSGW mice (Jackson Laboratory, Bar Harbor, ME, USA).

### Xenotransplantation assays

For primary transplantation of CD34+ cells electroporated with Cas9, RNP and RNP/GSE56, 25 mg/kg of busulfan (Busulfex, Otsuka, Rockville, MD, USA) was diluted with phosphate-buffered saline (PBS) to a final volume 200 uL and injected into twelve- to thirteen-week-old female NSG mice (Jackson Laboratory) intraperitoneally 24 hours before tail-vein injection. PB was sampled at indicated times post-transplant. For the endpoint analysis in primary transplant (22 weeks post-transplant), BM and spleen were also harvested. Cells from PB, BM and the spleen were assessed with the following antibody panel (all purchased from Becton Dickinson (BD), Franklin Lakes, NJ, USA): (1) Lineage staining (PB, BM and spleen samples): PE-anti-CD45, PE-Cy7-anti-CD20, APC-anti-CD13, APC-Cy7-anti-CD3 (all in 1:25 dilution); (2) HSC staining (BM samples only): PE-anti-CD45, FITC- or AlexaFlour700- anti- CD34, APC-anti-CD38 (all in 1:25 dilution). For purification of human CD45+ cells from primary transplanted mice, the remaining fresh murine BM cells were flow sorted and subsequently cryopreserved in CryoStor® CS5 (STEMCELL Technologies). For secondary transplant, 10 mg/kg of busulfan was diluted with PBS to a final volume 200 uL and injected into twelve- to thirteen-week-old female NBSGW mice (Jackson Laboratory) intraperitoneally 24 hours before tail-vein injection. Low dose busulfan conditioned NBSGW female mice were chosen for the limiting-dilution secondary transplantation since this xenograft model was previously shown to be more sensitive for detection of human cell engraftment than non-conditioned NBSGW mice or sublethally irradiated NSG mice (Jackson Laboratory) ^60, 61^. For the endpoint analysis in secondary transplant (20 weeks post-transplant), BM was harvested and assessed with the following antibody panel: PE-anti-CD45, PE-Cy7-anti-CD20, APC-anti-CD13 (all in 1:25 dilution). A secondary mouse was scored as positive if multilineage defined by clear evidence of myeloid (CD13+) and lymphoid (CD20+) engraftment and the percentage of human CD45+ cells was > 0.01%. The frequency of LT-HSCs was determined by an LDA, as previously reported^62^. The HSC frequency was calculated using ELDA software (http://bioinf.wehi.edu.au/software/elda/) and plotted using limdil function in satmod package within the RStudio IDE.

For primary transplantation of G-CSFR^hi^ CD34+ cells, FACS sorted cells were transplanted into non-conditioned twelve- to thirteen-week-old NBSGW female mice by tail-vein injection. For the endpoint analysis (16 weeks post-transplantation), cells from murine BM were assessed with the following antibody panels: (1) Lineage staining: PE-anti-CD45, PE-Cy7-anti- CD20, APC-anti-CD13, APC-Cy7-anti-CD3 (all in 1:25 dilution); (2) HSC staining: PE-anti- CD45, FITC- anti-CD34, APC-anti-CD38 for HSC (all in 1:25 dilution). For secondary transplantation, human CD45+ cells were purified from the remaining fresh murine BM cells using human CD45 MicroBeads (Miltenyi Biotec), and subsequently transplanted into twelve- to thirteen-week-old low-dose busulfan conditioned NBSGW female mice. For the endpoint analysis in secondary transplants (16 weeks post-transplant), BM was harvested and assessed with the following antibody panel: PE-anti-CD45, PE-Cy7-anti-CD20, APC-anti-CD13, APC- Cy7anti-CD3, FITC-anti-CD34 (all in 1: 25 dilution).

Animals were housed and handled in accordance with the guidelines set by the Committee on Care and Use of Laboratory Animals of the Institute of Laboratory Animal Resources, National Research Council (DHHS publication No. NIH 85-23), and the protocol was approved by the Animal Care and Use Committee of the National Heart Lung and Blood Institute.

### G-CSF administration and considerations

125 µg/kg of recombinant human G-CSF (Amgen) was diluted with PBS to a final volume 100-150 uL, and subcutaneously (SC) injected into each mouse once daily for the indicated duration.

There is no current consensus on the dose, frequency and duration of recombinant human G-CSF use in mice. In a previous study in humans, administration of 11.5 µg/kg SC resulted in a peak serum concentration of 49 ng/mL within 2 to 8 hours and an elimination half-life of approximately 3.5 hours (Amgen’s package insert). In mice, an early study observed a similar half-life (3.1 hours) after injecting 100 µg/kg SC into C57Bl/6x129SvJ mice^63^. Another study showed a peak plasma concentration of approximately 50-100 ng/mL after injecting 10 µg SC into 8 week–old female outbred Hsd:ICR mice^64^. Assuming a ∼20 g mouse (the approximate weight of 10–12-week C57Bl/6J female mouse), the dose used in Scholz and colleagues’ study was approximately 500 µg/kg SC. Finally, the conversion of the recommended 10 μg/kg/day SC for human patients was estimated as ∼125 μg/kg/day SC in mice using a surface area conversion from human to mouse^65, 66^. After considering the expected peak serum concentration and elimination half-life of recombinant human G-CSF in mice and surface area conversion from human to mouse suggested by aforementioned studies, we chose the conventional once-daily dose of G-CSF (125 ug/kg/day SC) for our *in vivo* experiments.

### DNA isolation and T7 Endonuclease I Assay

Genomic DNA was isolated from bulk HSPC samples (∼2 x 10^6^ cells per sample) using a QIAGEN Blood & Tissue DNA isolation kit (QIAGEN, Inc., Germantown, MD, USA) following the manufacturer’s protocol. For flow sorted human CD45+ cells, genomic DNA was extracted using a NucleoSpin Tissue XS kit (TAKARA Bio USA) following the manufacturer’s protocol. Cas9 target sites within the AAVS1 locus were PCR amplified in a 25 or 50 μL reaction volume per sample using high-performance RANGER PCR Mix (Bioline USA, Inc., Memphis, TN, USA) or PlatinumTM SuperFiTM PCR Master Mix (Thermo Fisher Scientific) and ∼5 ng of genomic DNA and 100 ng each of the appropriate forward and reverse primer. Target-specific primer sets were: AAVS1-F, 5’-CTTGCTTTCTTTGCCTGGAC-3’; AAVS1-R, 5’-ACACCTAGGACGCACCATTC-3’. PCR parameters were as follows: 98°C for 3 min, followed by 35 rounds of 98°C for 15 s, 60°C for 30 s, 72°C for 2 min, with a final extension step at 72°C for 5 min. After amplification, each PCR reaction was partially purified using a BioRad PCR Kleen column (BioRad Laboratories, Hercules, CA, USA) or NucleoSpin® Gel and PCR Clean- up (Takara Bio USA) according to the manufacturer’s instructions. T7 endonuclease I (T7E1) reactions were assembled in 8-well, 0.2 mL PCR strips with 20-200 ng of column-purified PCR product per reaction. Amplicons were heat-denatured and re-annealed in a programmable thermocycler using the following hybridization conditions: 95°C for 5 min, ramp from 95 to 85°C at -2°C per second, ramp from 85 to 25°C at -0.1°C per second, followed by a “hold” at 22°C. After re-annealing, 10 units of T7E1 (New England Biolabs, Inc., Ipswich, MA) were added to the appropriate tubes, and the samples were incubated at 37°C for 15 min. After T7E1 digestion, 4 μL of NOVEX. 5X Tris-borate-EDTA (TBE) gel loading buffer was directly added to each PCR tube. A portion of each sample was loaded onto a pre-cast Novex 4–20% polyacrylamide TBE gel (Thermo Fisher Scientific), followed by electrophoresis at 150 volts (constant voltage) until the lower tracking dye approached the bottom of the gel. The gels were stained using SYBR Green I nucleic acid gel stain (Thermo Fisher Scientific) and visualized using a Benchtop 3UV transilluminator (UVP). Densitometric quantification of DNA bands was done using ImageJ software. Indel frequencies were calculated using the formula, where the fraction cleaved = ([density of digested products] / [density of digested products + density of undigested parental band]) as described previously^67^. Estimated Indel frequencies > 0.1 using this formula were considered for analyses of Indel frequencies within human CD45+ cells isolated from primary mice comparing G-CSF treated vs. untreated groups.

### Lentiviral vector preparation

The χHIV vector was prepared by four-plasmid co-transfection of 293T cells with χHIV Gag/Pol, HIV1 Rev/Tat, VSV-G envelope, and GFP-expressing SIN-HIV1 vector plasmids, as previously described^68^. In all vectors, the transgenes were driven by an EF1α promoter. Biological titers of lentiviral vectors were measured in human CD34+ cells.

### Transduction of human CD34+ HSPCs

Human CD34+ HSPCs were cultured overnight on RetroNectin-coated (Takara Bio USA) plates in StemSpan SFEM II medium supplemented with SCF 100 ng/mL, TPO 100 ng/mL, FLT3 100 ng/mL and 1% human albumin, and transduced with GFP-expressing lentiviral vectors in the presence of 4 μg/ml protamine sulfate (Sigma-Aldrich, St. Louis, MO, USA).

### Quantitative reverse transcription PCR

RNA was extracted using the RNeasy Plus Mini Kit (Qiagen) according to the manufacturer’s instructions. The genomic DNA elimination columns contained in the kit were used to eliminate possible DNA contamination during the extraction. Quantitative reverse transcription (RT) and real-time PCR (qRT-PCR) were done using TaqMan™ RNA-to_Ct 1-Step kit (Thermo Fisher Scientific, AB 4392938) on BioRad CFX 96 Real-time systems with the following TaqManTM probes and primers purchased from Thermo Fisher Scientific; ACTB-VIC (Hs01060665_g1) and CDKN1A-FAM (Hs00355782_m1).

### Confocal microscopy

Cover glasses (No. 1.5, 24 x 60 mm) (Fisher Scientific, Hampton, NH, USA) were treated with Poly-D-lysine solution (Sigma-Aldrich) at 0.1 mg/ml concentration for 60 minutes at room temperature (RT). After three washes with PBS, approximately 0.5-1x10^5^ cells were seeded on coated cover glasses for 30 minutes and then fixed with chilled 4% paraformaldehyde (Santa Cruz Biotechnology, Dallas, TX, USA) for 10 minutes at RT. After three washes with PBS, cells were then permeabilized with 0.1% Triton X-100 for 10 minutes and blocked with 5% BSA for 30 minutes at RT. Cells were then stained with rabbit anti-53BP1 antibody (Bethyl Laboratories, Montgomery, TX, USA, A700-011) and mouse anti-phospho-histone H2A.X (Ser139) antibody (MilliporeSigma, Burlington, MA, USA, 05-636) in 1:200 dilution at 4°C overnight. The next day (after staining with primary antibody), cells were washed three times with PBS and stained with anti-rabbit IgG antibody conjugated to Alexa488 (Sigma-Aldrich, SAB4600045) and anti-mouse IgG antibody conjugated to Alexa647 (Thermo Fischer Scientific, A21235) in 1:200 dilution at RT for 60 minutes. After staining with secondary antibody, cells were washed three times with PBS, and cover glasses were then mounted with ProLong™ Gold Antifade Mountant with DAPI (Thermo Fisher Scientific) on glass slides. Fluorescent images were acquired using a Leica SP8 confocal microscope (Leica Microsystems, Mannheim, Germany) in the upright configuration using a Leica 63x (1.4NA) HC PL APO objective lens and standard (DAPI) or hybrid (Alexa488 and Alexa647) PMTs. Tiled z-stack volumetric images of DAPI, Alexa488, and Alexa 647 were acquired sequentially with 405 nm, 488 nm, and 638 nm laser excitation and 412-490 nm, 500-600 nm, and 650-750 nm bandpass emission collection, respectively. All images were taken at a speed of 600 Hz, a pinhole setting of 1 AU, pixel sizes of 345 nm in the XY plane, interslice spacing of 500 nm, and line average of two. The collected 3D tiled images were merged using the Leica LAS X software (version 3.5.7). Quantification of 53BP1 foci in immunofluorescence images was conducted using the 3D cell analysis machine learning algorithm in Aivia software (version, 12.0.0, Leica Microsystems).

### CITE-sequencing

The 3’ end scRNA-seq was performed on a Chromium Single-Cell Controller (10X genomics, Pleasanton, CA, USA) using the chromium Single Cell 3’ Reagent Kit v3.1 dual index kit according to the manufacturer’s instructions. Human CD34+ cells were electroporated with Cas9 alone or AAVS1-specific RNP in the presence or absence of GSE56 mRNA and subsequently transplanted into busulfan conditioned NSG mice. At 24 hours post-transplantation, transplanted mice started receiving daily G-CSF or PBS. After 3 days of daily G-CSF or PBS injection (at 96 hours post-transplantation), mice were euthanized, and BM cells were harvested. Human CD34+ cells were purified from fresh BM cells using MicroBead kit (Miltenyi Biotech 130-046-702) and subsequently cryopreserved in CryoStor® CS5 (STEMCELL Technologies).

On the day of library preparations, frozen cells were thawed using chilled IMDM (Thermo Fisher Scientific) supplemented with 2% fetal bovine serum (FBS), washed twice, and counted with AOPI staining on an automated cell counter. Between 50,000 to 100,000 cells were resuspended in 10 uL of PBS supplemented with 2% FBS and 2 mM EDTA (hereafter named “FACS buffer”) together with 5 uL of TruStain FcX (BioLegend, San Diego, CA, USA). Cells were incubated with the FcX block for 10 minutes at RT, after which 1 uL each of CD45-PE (BD) and CD34-FITC (BD) as well as 0.5 ug each of the 10 TotalSeq-B oligo-tagged antibodies (BioLegend) listed in Supplementary Table 3 were added and cell pellets were suspended. After incubation with these antibodies on ice for 30 minutes, cells were washed three times. Once 1 uL of 7-AAD (Thermo Fisher Scientific) was added, cells were filtered through 40 um cell strainers and fluorescence-activated cell sorting (FACS) was performed to sort 2,500 live single CD45+CD34+ cells from each experimental group. Immediately after sorting, all cells from each experimental group were pooled and centrifuged at 3000 rpm for 30 minutes using a microcentrifuge, and the supernatant was aspirated. All recovered cells were utilized for the subsequent procedure (estimated recovery: 2,000-2,500 cells/ per experimental group).

Oligo-tagged cells were loaded on a Chromium Chip B with master mix, gel beads, and partitioning oil, again according to 10x Genomics instructions. 3’ v3.1 GEX libraries were constructed according to 10x Genomics protocol. 3’ v3.1 GEX and ADT libraries were pooled in a 2:1 ratio and sequenced with an Illumina NovaSeq SP flow cell (100 cycles, 2 lanes). Target sequencing depth for the GEX libraries was 20,000 read pairs per cell and 5,000 read pairs per cell for the ADT libraries.

Illumina sequencer’s base call files (BCLs) were demultiplexed, for each flow cell directory, into FASTQ files using Cellranger mkfastq with default parameters (v 6.1.2) in NIH Helix/Biowulf High Performance Computation (HPC) system. FASTQ files were then processed using Cellranger count with default parameters in NIH Helix/Biowulf HPC system. Internally, the software relies on STAR for aligning reads to a pre-build filtered human reference genome relying on GRCh38, while genes are quantified using ENSEMBL genes as gene model. The output of Cellranger is a filtered gene-barcode matrix containing the UMI counts for each gene.

Gene counts were processed with Seurat (v 4.1.0, https://satijalab.org/seurat/). Quality control was performed to filter low-quality cells by excluding cells expressing genes more than [mean + 2 x standard deviations (SD)] of total genes, less than [mean - 2 x SD] of total genes, or mitochondrial genes comprising more than 10% of the total gene expression per cell. Counts were normalized using Seurat function NormalizeData with default parameters. Expression data were then scaled using the ScaleData function. The HTODemux function was used to demultiplex cells to their samples of origins and identify doublets and negative cells for further filtering. To decrease noise and make downstream computations more tractable, the dataset was reduced to 20 dimensions of PCA using the RunPCA function. For visualization, the dataset was dimensionally reduced to two dimensions of Uniform Manifold Approximation and Projection (UMAP) using RunUMAP function. Shared-nearest neighbor (SNN) graph based on cell-cell distance matrix constructed using FindNeighbors function was subsequently used to partition cells into clusters using Louvain modularity optimization algorithm using FindClusters function. This resulted in 7 distinct HSC and progenitor clusters.

HSC and progenitor clusters were first annotated by computing module scores of previously reported HSPC gene set^69–71^ using a previously described approach^72, 73^. We assigned each cluster a specific HSPC identity by statistically comparing module scores and assigning the top-scoring cell type to each cluster. In this manner, we annotated seven clusters, including CMP, CLP, LMPP, GMP, MLP, MEP, and HSC clusters. We next confirmed expression levels of lineage defining genes within these clusters: Musashi-2 (MSI2) and Hepatic leukemia factor (HLF)^74–81^ expression in the HSC cluster; Myeloperoxidase (MPO)^82^ expression in the GMP cluster; GATA2^83^ expression in the MEP cluster, and CD52^84^ expression in the CLP cluster. The CMP cluster was annotated by being enriched in the progenitor libraries and being positioned within the megakaryocyte-erythroid branch by pseudotime analysis, but lacking expression of other specific lineage markers as defined. The LMPP cluster was annotated by being enriched in the progenitor (most notably lymphoid) libraries and being positioned between HSC cluster (designated as pseudotime origin) and myeloid/lymphoid branches by pseudotime analysis, but lacking expression of other specific lineage markers as defined. Surface protein expression from CITE-seq data further confirmed that the transcriptionally defined HSC cluster 6 is consistent with the known HSC immunophenotype in humans, as evidenced by higher expression of CD90 and CD49f and lower expression of CD38 and CD45RA.

Pseudotime tools estimate the order of individual cells according to their progress through a biological process. Slingshot (v2.4) was used to create the sample pseudotime trajectories. The slingshot function of the slingshot package^85^ was run on the Seurat generated cell clusters with cluster 6 as the starting cluster. Each predicted path was then separately plotted using the UMAP generated in Seurat with the cells colored based on the estimated pseudotime.

Genes differentially expressed across different conditions were identified using the FindMarkers function, applying the MAST test^86^ and the Bonferroni correction. Average log_2_FC were computed after adding pseudocount of 0.001 to averaged expression values and only genes expressed in at least 1% of cells in at least one contrasting group were considered.

We used GSEA (v4.2.3) software from the GSEA Broad Institute website to carry out gene-set enrichment analysis on Hallmark gene sets, available in the Broad Institute Molecular Signatures Database^87^. Genes were ranked based on the direction of the fold change [±1 × −(log10(pvalue))] calculated by MAST test. The minimum and maximum number of genes in each pathway allowed for the GSEA was set to 5 and 500 respectively. FDR q-values were estimated to correct the p-values for testing multiple pathways.

### Cell cycle analysis and apoptosis assay

25 mg/kg of busulfan (Busulfex, Otsuka) was diluted with phosphate-buffered saline (PBS) to a final volume 200 uL and injected into twelve- to thirteen-week-old male NBSGW mice (Jackson Laboratory) intraperitoneally 24 hours before tail-vein injection. Pre-cultured human CD34+ cells were electroporated with Cas9 alone or AAVS1-specific RNP in the presence or absence of GSE56 mRNA and subsequently transplanted into busulfan-conditioned male NBSGW mice. At 24 hours post-transplantation, transplanted mice started receiving daily G-CSF or PBS. After 3 daily injections of G-CSF or PBS (96 hours post-transplantation), mice were euthanized, and BM cells were harvested.

Cell cycle status was analyzed using FITC BrdU Flow Kit (BD, 559619) according to the manufacturer’s instructions. Briefly, 3 x 10^6^ BM cells were incubated with human Fc BlockTM (BD, 564219) for 10 minutes at RT. Then, cells were stained with PE-anti-CD45, Alexaflour700- anti-CD34 and APC-anti-CD38 (all purchased from BD and used in 1:25 dilution) for 30 minutes on ice. After washing with FACS buffer three times, cells were fixed and permeabilized according to the manufacturer’s instructions. For BrdU labeling, a single dose of BrdU (1Lmg) was administered into each mouse intraperitoneally 24 hours prior to harvesting BM cells. After permeabilization, cells were treated with DNase to expose BrdU epitopes, and then stained with 1 uL of FITC-anti-BrdU antibody in 50 uL of BD Perm/Wash buffer for 20 minutes at RT. After washing with BD Perm/Wash buffer, cells were suspended in 500 uL BD Perm/Wash buffer containing 10 uL of the 7AAD solution and incubated for 15 minutes at RT. Then, cells were immediately analyzed using an LSR II Fortessa flow cytometer (BD), run at a rate no greater than 3,000 events per second. At minimum, a total of 2,000,000 events were acquired in each sample.

Cellular survival and apoptosis were quantified using Annexin V Apoptosis Detection kit (Thermo Fisher Scientific, 88-8006-74) according to the manufacturer’s instructions. Briefly, 1-2 x 10^6^ BM cells were incubated with human Fc BlockTM (BD, 564219) for 10 minutes at RT. Then, cells were stained with PE-anti-human CD45, Alexaflour700-anti-CD34, APC-anti-CD38 (all purchased from BD and used in 1:25 dilution) for 30 minutes on ice. After washing with 1X binding buffer twice, cells were stained using eFlour450-Annexin V for 15 minutes at RT. After cells were washed and resuspended in 1X binding buffer, 7-AAD viability staining solution (Thermo Fisher Scientific, 00-6993) was added, and stained cells were immediately analyzed using an LSR II Fortessa flow cytometer (BD). At minimum, a total of 1,000,000 events were acquired in each sample.

### Statistics

Data are reported as mean ± standard error of the mean (SEM). Statistical significance (p < 0.05) was assessed using two-sided unpaired Student t-test, with Bonferroni multiple comparison test as indicated, or one-way/two-way ANOVA or Kruskal-Wallis test with Tukey correction or Sidak multiple comparison test as indicated. Statistical analyses for bulk and single cell RNA sequencing experiments were performed within the RStudio IDE. All the other statistical analyses comparing the different experimental groups were performed using GraphPad Prism 9 software version 9.5.1.

## Data availability

CITE-sequencing data are available at dbGap via authorized access request as detailed at https://dbgap.ncbi.nlm.nih.gov/aa/wga.cgi?page=login (dbGap accession number: phs003277.v1.p1.). Data are also available in Gene Expression Omnibus under accession GSE230182.

## Acknowledgements

The authors thank all members of the Larochelle Lab for technical support and helpful discussions; Suk See De Ravin for sharing GSE56 mRNA; David Stroncek and the NIH Department of Transfusion Medicine and Cell Processing Section staff for apheresis, selection and cryopreservation of human CD34+ cells; Richard Gustafson and the outpatient clinic nursing staff for recruiting normal volunteers and providing G-CSF administration teaching to healthy subjects; Temeri Wilder-Kofie, James Hawkins and Building 50 Animal Facility staff for excellent animal care. Figures 1a, 1f, 2a, 3a, 4a, 5a, 5f and the graphical abstract were designed using BioRender.com. This work was supported by the Intramural Research Program of the National Heart, Lung, and Blood Institute, National Institutes of Health, USA (ZIA HL006172 and Z99 HL999999).

## Author contributions

D.A. and A.L. conceptualized the project and designed the experiments. D.A. performed all experiments and acquired all data. D.A., Y.Lu., P.L., and Y.Li. prepared CITE-seq libraries. D.A., V.C. and N.R. analyzed CITE-seq datasets. D.A. and C.S.R. harvested and processed murine bone marrow and spleen samples for flow cytometry analysis. D.A. and P.D. performed fluorescence-activated cell sorting. R.H.S. designed sgRNA for gene editing and primers for PCR reactions and helped with T7I assay. D.A. and C.C. acquired and analyzed confocal microscopy data. D.A. and A.L. wrote the original draft. All authors have read and agreed to the published version of the manuscript.

---

**Extended Data Fig. 1.**
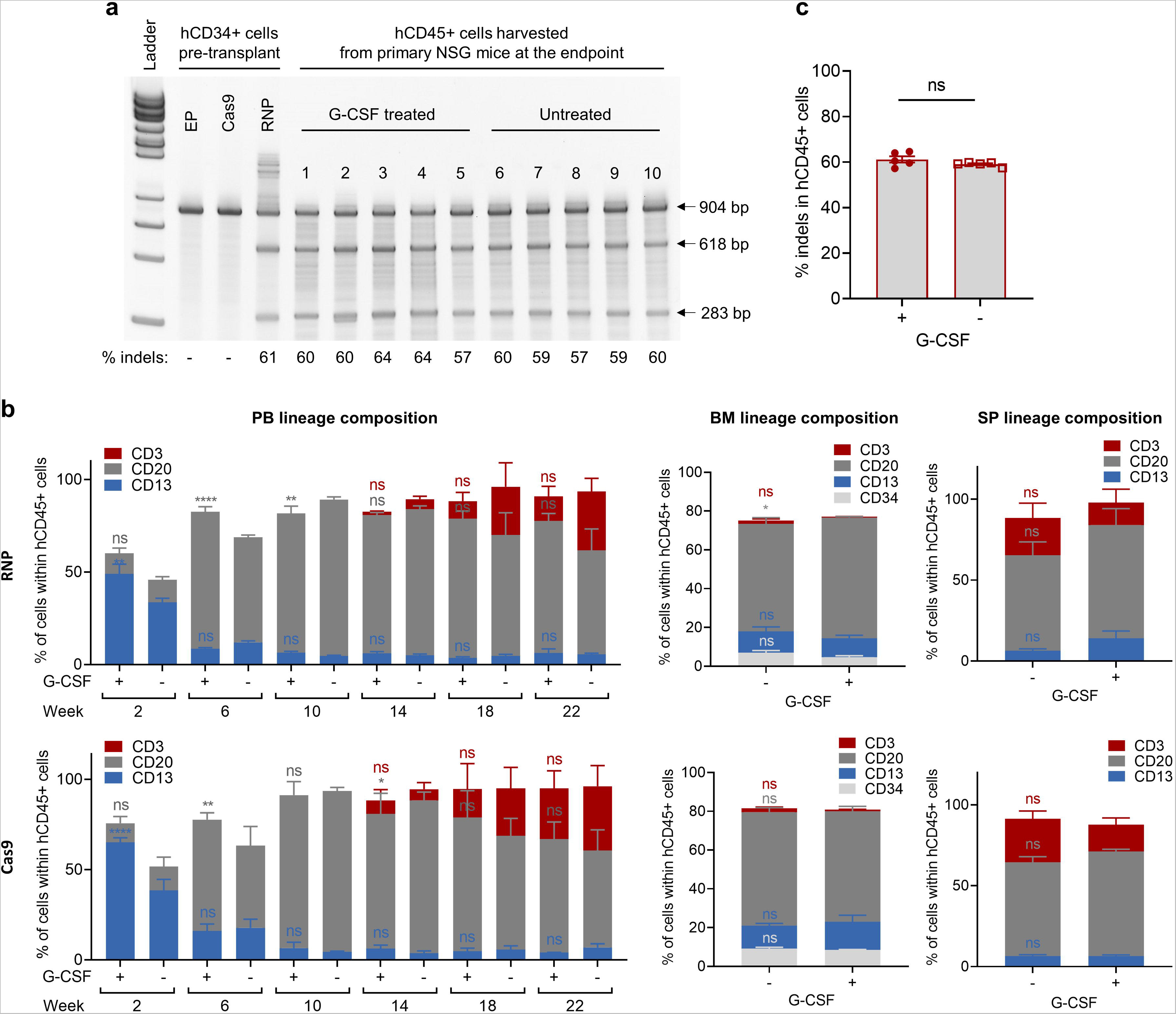

**Extended Data Fig. 2.**
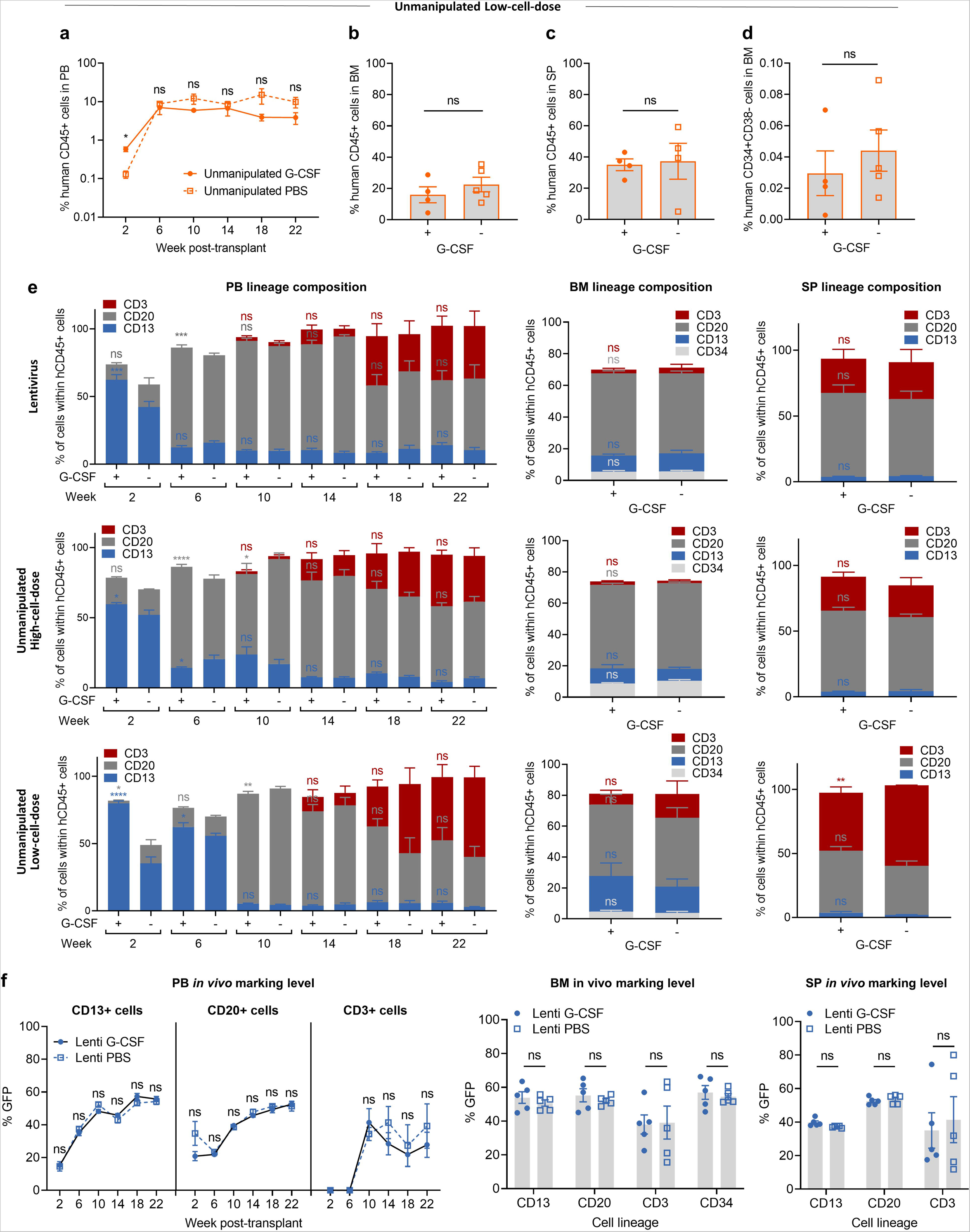

**Extended Data Fig. 3.**
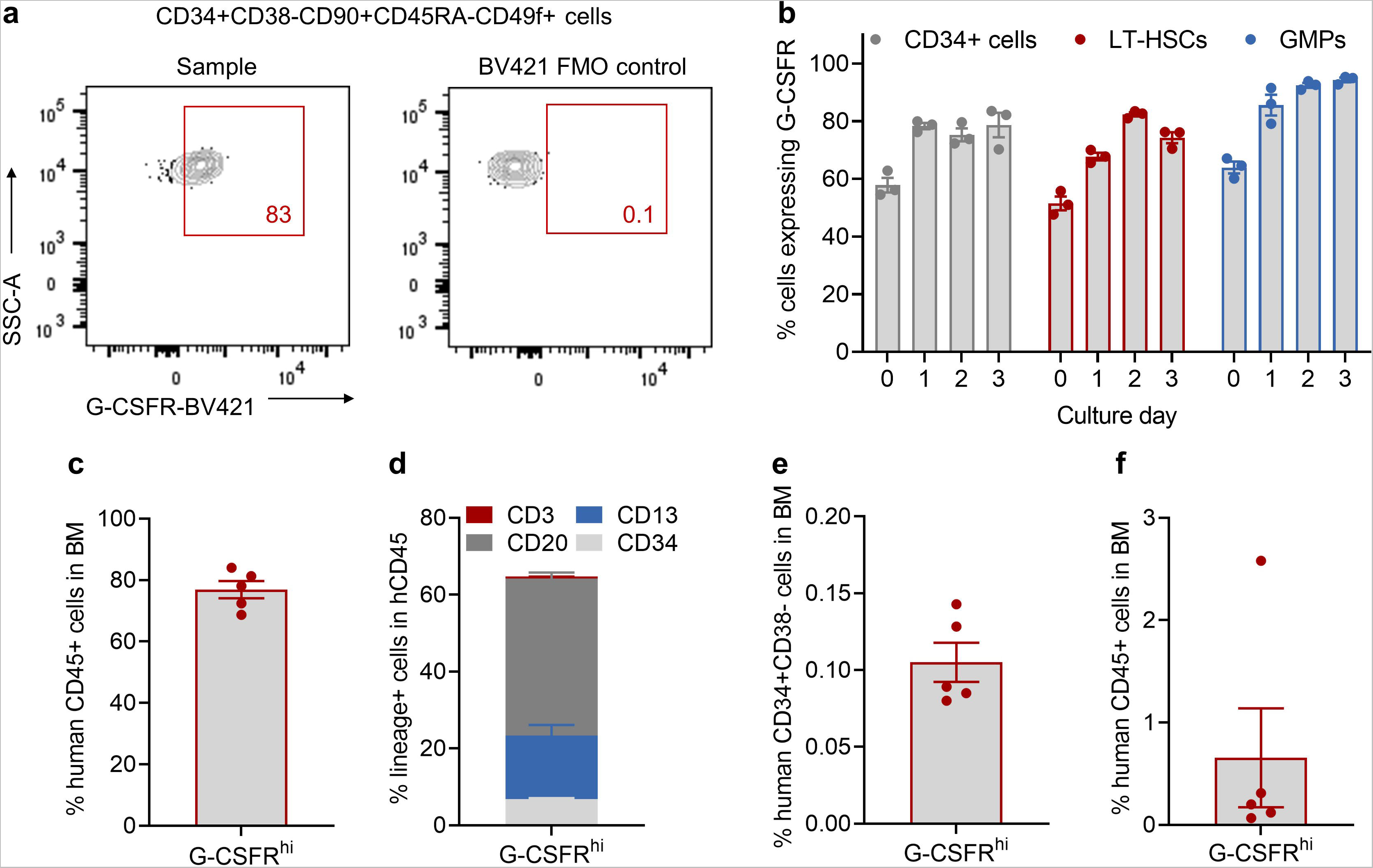

**Extended Data Fig. 4.**
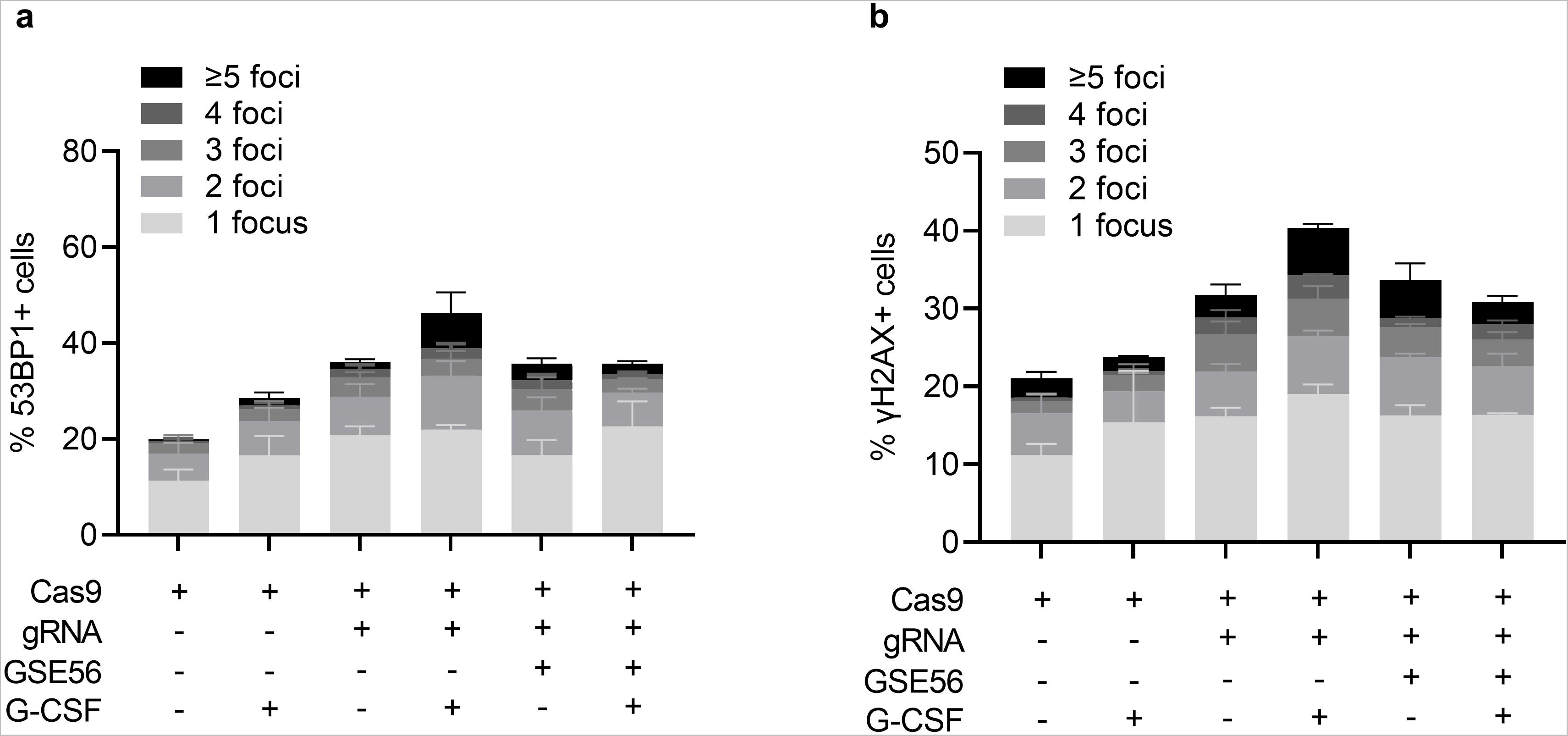

**Extended Data Fig. 5.**
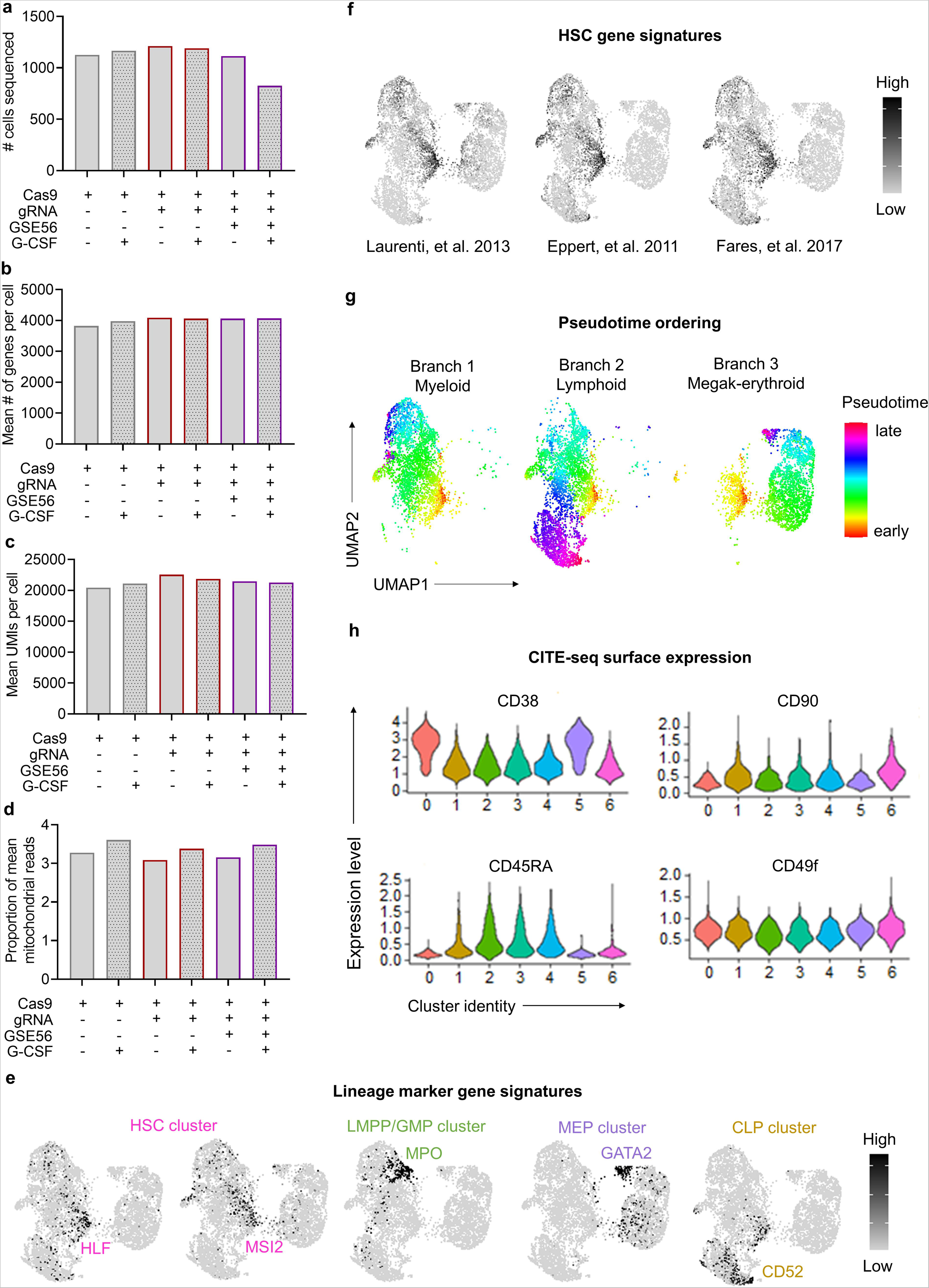

**Extended Data Fig. 6.**
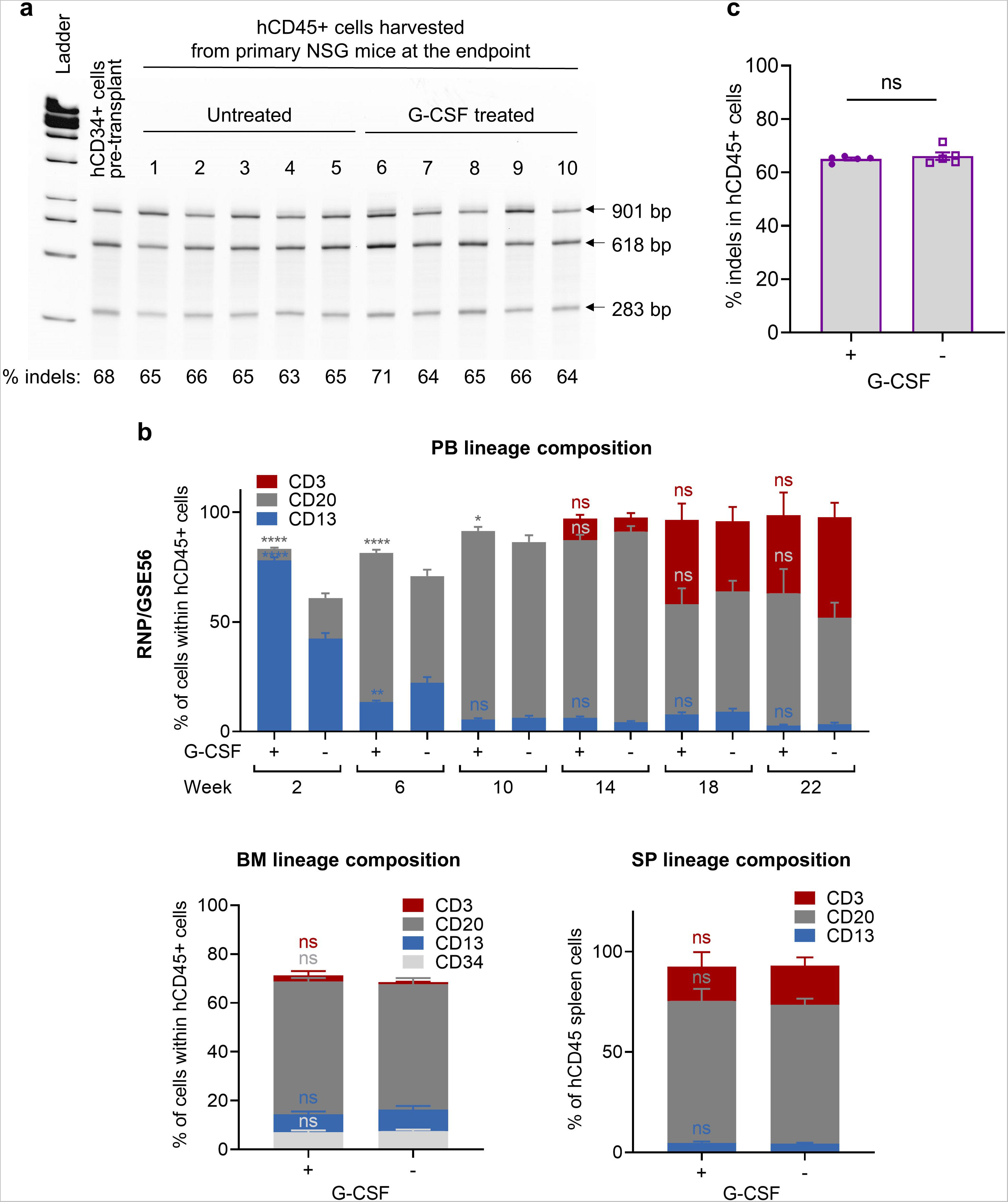

