## Supplemental Material for "Post-Transplant Administration of G-CSF Impedes Engraftment of Gene Edited Human Hematopoietic Stem Cells by Exacerbating the p53-Mediated DNA Damage Response"

**Affiliations:**

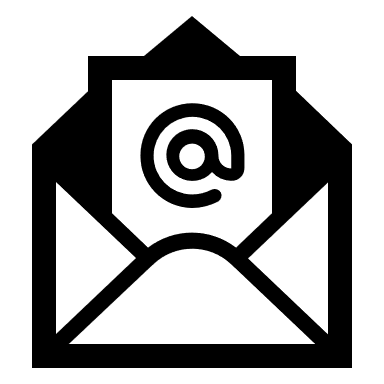

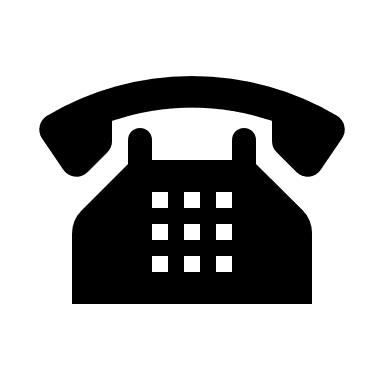

(301) 814-8289

This file includes:

- **Supplementary Figures**
- **Extended Data Fig. 1:** Analysis of gene editing frequency and lineage composition within human CD45+ cells after primary transplantation in the RNP and Cas9 groups.
- **Extended Data Fig. 2**: Post-transplant G-CSF administration has minimal impact on long-term engraftment of lentiviral vector transduced or unmanipulated HSPCs.
- **Extended Data Fig. 3:** The G-CSF receptor is expressed on human HSPCs and HSPCs bearing the G-CSF receptor can sustain long-term multilineage hematopoietic reconstitution after transplantation.
- **Extended Data Fig. 4:** G-CSF exacerbates the p53 DDR activated by Cas9-induced DNA DSBs and transient p53 inhibition mitigates the G-CSF effects in human HSPCs.
- **Extended Data Fig. 5:** Cellular Indexing of Transcriptomics and Epitopes sequencing (CITE-seq) analysis of Cas9-, RNP- and RNP/GSE56-electroporated human HSPCs.
- **Extended Data Fig. 6:** Analysis of gene editing frequency and lineage composition within human CD45+ cells after primary transplantation in the RNP/GSE56 group.

Supplementary Tables are provided in a separate Excel document:

- **Supplementary Table 1**: Complete list of differentially expressed genes within the HSC cluster of human CD34+ cells analyzed by CITE-sequencing using the MAST test.
- **Supplementary Table 2**: Complete list of upregulated and downregulated hallmark gene sets in human CD34+ cells analyzed by CITE sequencing.
- **Supplementary Table 3**: Key Resources Table.

Extended Data Fig. 1

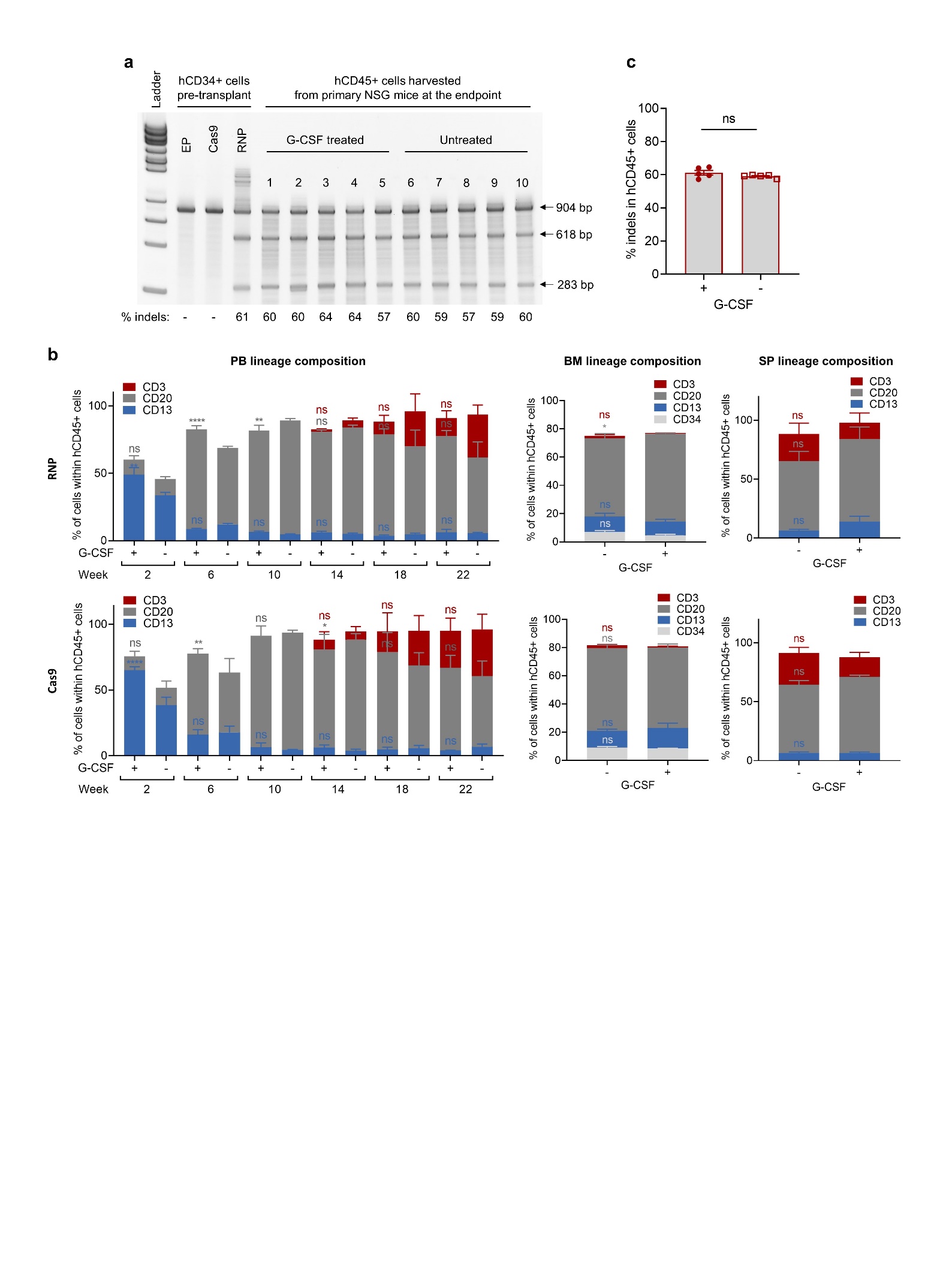

**Extended Data Fig. 1 | Analysis of gene editing frequency and lineage composition within human CD45+ cells after primary transplantation in the RNP and Cas9 groups. a**, Gel image of the T7 endonuclease I (T7EI) assay for quantification of sgRNA/Cas9 RNP-mediated indels after electroporation (EP) in human CD34+ (hCD34+) cells pre-transplant or human CD45+ (hCD45+) cells isolated from murine bone marrow (BM) after primary transplantation. **b**, Lineage composition (CD3+ T lymphoid, CD20+ B lymphoid, CD13+ myeloid and CD34+ progenitor cells) within hCD45+ cells in the peripheral blood (PB), BM or spleen (SP) of NSG mice after primary transplantation in the RNP and Cas9 groups. **c**, Summary of indel frequencies within hCD45+ cells harvested from the BM of NSG mice after primary transplantation as measured by T7EI assay. In all panels, data are displayed as mean ± standard error of the mean (SEM). Two-way ANOVA with Sidak multiple comparison test was used in panel b, and two-sided unpaired t-test was used in panel c. ns, not significant, * p ≤ 0.05, ** p ≤ 0.01, **** p ≤ 0.0001. Associated with Fig. 1.

Extended Data Fig. 2

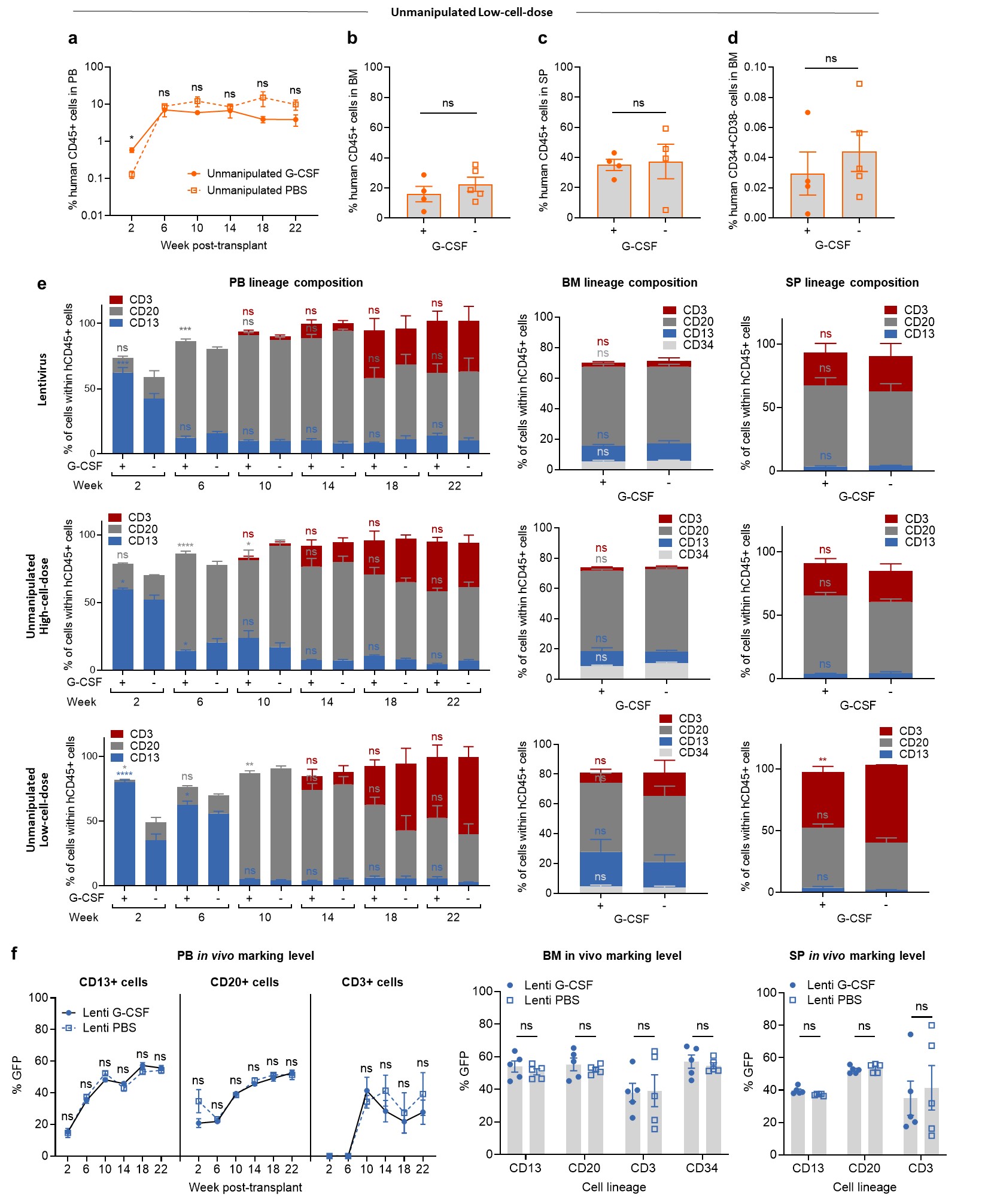

**Extended Data Fig. 2 | Post-transplant G-CSF administration has minimal impact on long-term engraftment of lentiviral vector transduced or unmanipulated HSPCs. a-d,** A total of 1 x 10^5^ unmanipulated human mobilized peripheral blood (MPB) CD34+ cells were transplanted into NSG mice after busulfan conditioning. G-CSF or PBS was subcutaneously injected from post-transplant day 1 to 14 and hematopoietic reconstitution was compared between both groups (n = 4-5 mice/group). Percentages of human CD45+ cells in the peripheral blood (PB) (**a**), bone marrow (BM) (**b**) and spleen (SP) (**c**), and percentages of HSC enriched populations (CD34+CD38- cells) in the BM (**d**) are shown. **e**, Lineage composition within human CD45+ cells in the PB, BM and SP after primary transplantation in the lentivirus, unmanipulated high-cell-dose and unmanipulated low-cell-dose groups. **f**, Percentages of GFP+ cells within each restricted cell lineage in the PB, BM or SP in the lentivirus group. In all panels, data are displayed as mean ± standard error of the mean (SEM). Two-way ANOVA with Sidak multiple comparison test was used in panel e. Otherwise, two-sided unpaired t-test was used. ns, not significant, *** p ≤ 0.001, **** p ≤ 0.0001. Associated with Fig. 1.

Extended Data Fig. 3
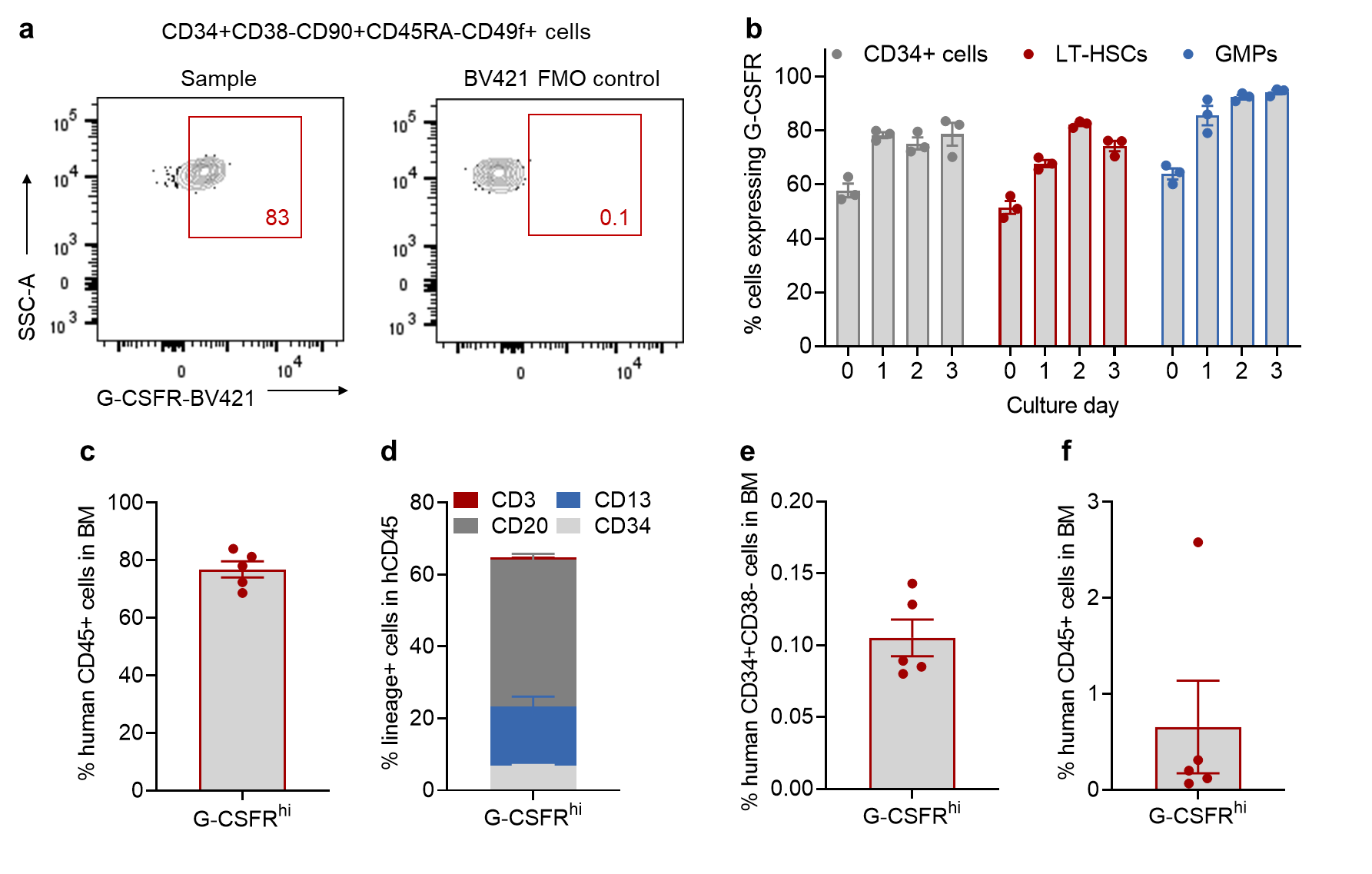

**Extended Data Fig. 3 | The G-CSF receptor is expressed on human HSPCs and HSPCs bearing the G-CSF receptor can sustain long-term multilineage hematopoietic reconstitution after transplantation.** Surface G-CSF receptor (G-CSFR) expression was measured by flow cytometry on human mobilized peripheral blood (MPB) CD34+ HSPCs before or after up to 3 days of culture to mirror culture condition and duration utilized in Fig. 3a. To confirm the long-term repopulating function of CD34+ cells bearing the G-CSF receptor, we used fluorescence-activated cell sorting to select G-CSFR^hi^ (top 10%) HSPCs and transplanted sorted cells into immunodeficient non-obese diabetic (NOD)-B6-severe combined immunodeficiency (SCID)-IL2Rg^null^-^KitW41/W41^ (NBSGW) mice (50,000 cells/mouse). Human CD45 chimerism was assessed within the bone marrow (BM) at 16 weeks post-transplantation. To gain insights into the self-renewal potential of G-CSFR^hi^ HSPCs, human CD45+ cells isolated from the BM of primary mice were injected into secondary NBSGW mice and BM engraftment was analyzed 16 weeks following secondary transplantation. **a**, Representative flow cytometry profiles of MPB CD34+ cells highly enriched in long-term repopulating potential (LT-HSCs: CD34+CD38-CD90+CD45RA-CD49f+) after 2 days in culture. A Fluorescence Minus One (FMO) control sample was used to set a positive boundary for G-CSFR expression within LT-HSCs. **b**, Percentages of bulk CD34+ cells, LT-HSCs and granulocyte-monocyte progenitors (GMPs: CD34+CD38+CD45RA+) expressing G-CSFR (n=3 independent donors). **c**, Percentages of human CD45+ cells in the BM after primary transplantation of G-CSFR^hi^ CD34+ cells into immunodeficient NBSGW mice (n = 5 mice/group). **d**, Lineage composition within human CD45+ cells in the BM after primary transplantation. **e**, Percentages of LT-HSC enriched populations (CD34+CD38-) within the BM of primary mice (n = 5 mice/group). **f**, Percentages of human CD45+ cells in the BM after secondary transplantation (n = 5 mice/group). In all panels, data are displayed as mean ± standard error of the mean (SEM).

Extended Data Fig. 4
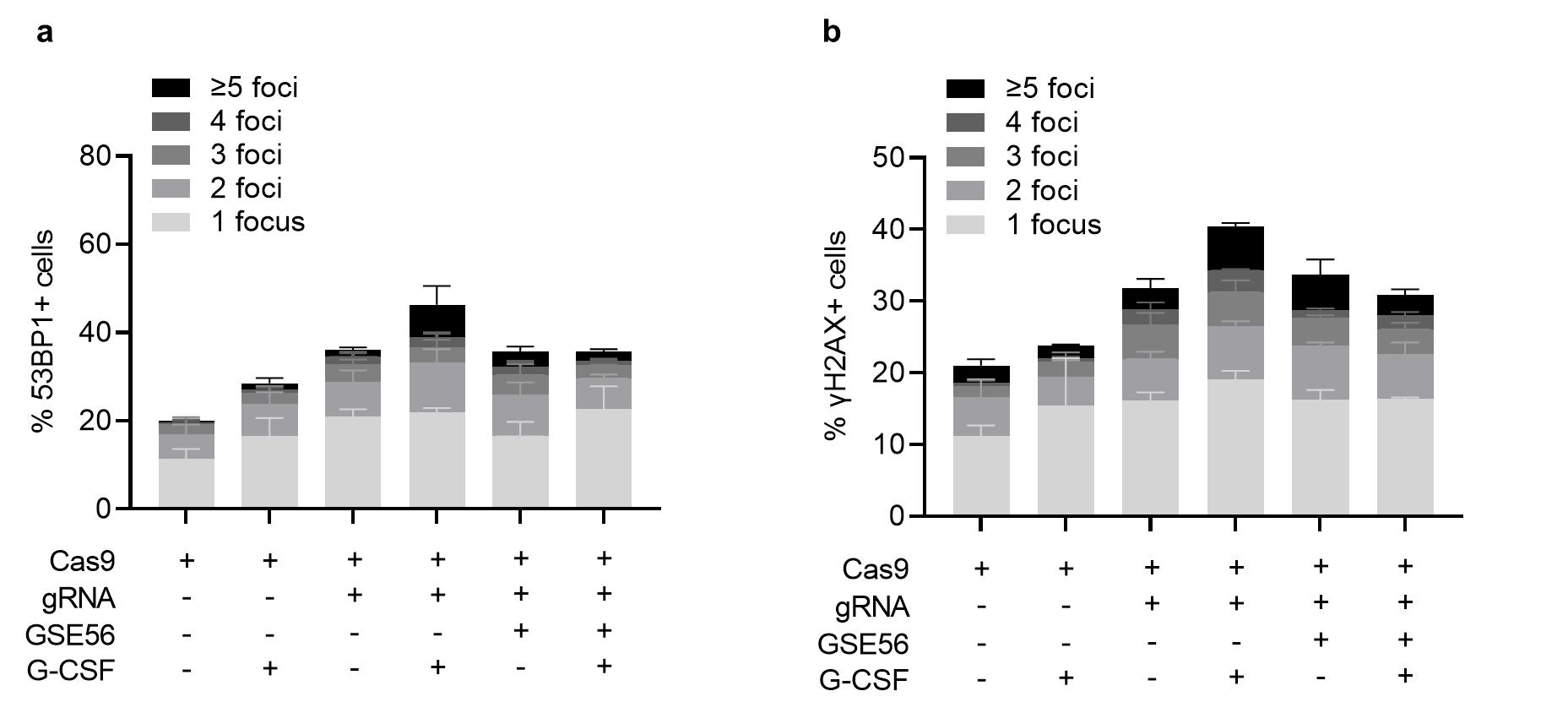

**Extended Data Fig. 4 | G-CSF exacerbates the p53 DDR activated by Cas9-induced DNA DSBs and transient p53 inhibition mitigates the G-CSF effects in human HSPCs.** Microscopically visible 53BP1 and γH2AX subnuclear foci were quantified in Cas9-, RNP- and RNP/GSE56-electroporated human CD34+ cells cultured with and without G-CSF as outlined in Fig. 3a. **a**, Percentages of 53BP1 foci-bearing cells (n = 3 independent donors/group). **b**, Percentages of γH2AX foci-bearing cells (n = 3 independent donors/group). Data are displayed as mean ± standard error of the mean (SEM). Associated with Fig. 3.

Extended Data Fig. 5
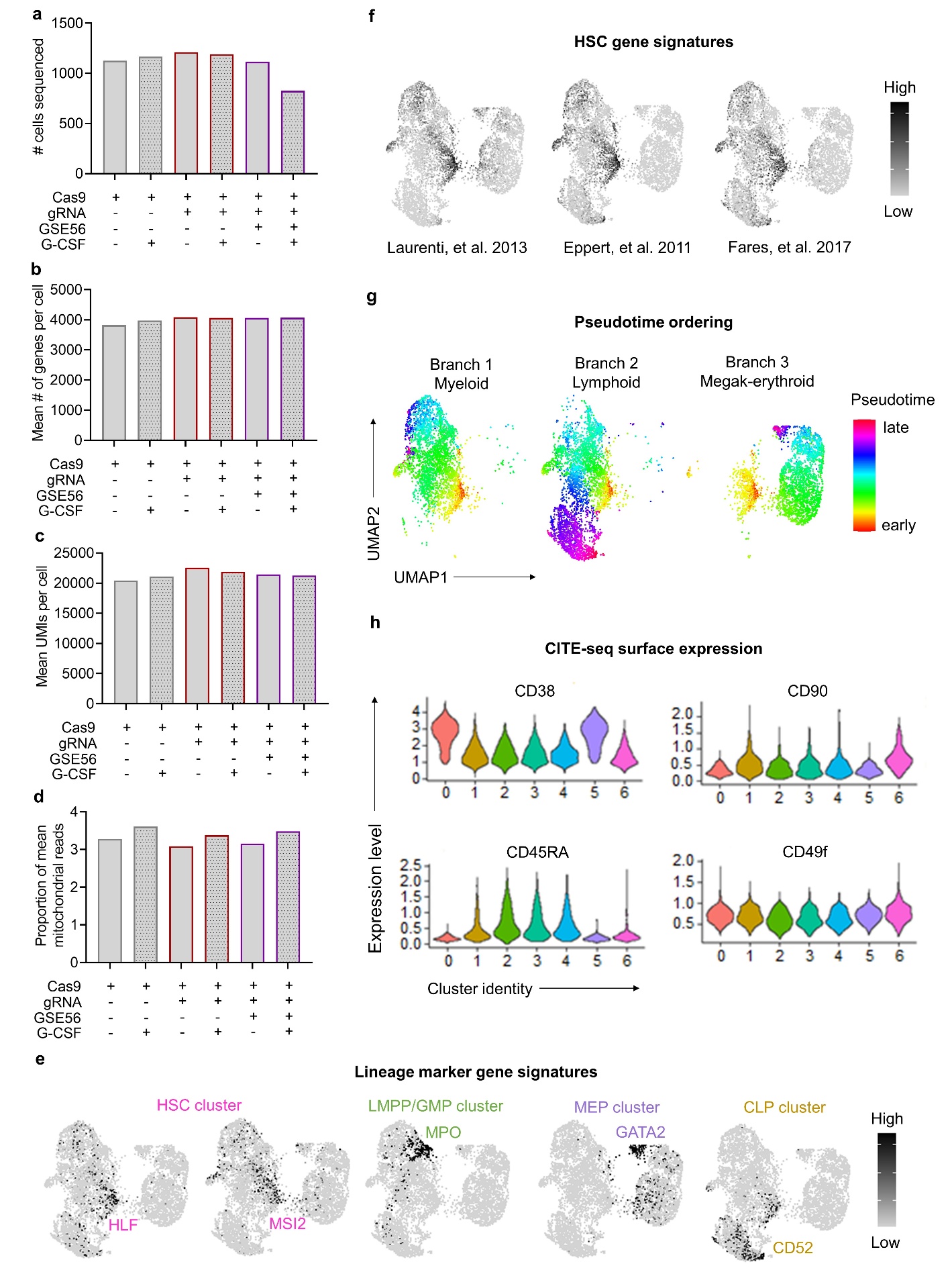

**Extended Data Fig. 5 | Cellular Indexing of Transcriptomics and Epitopes sequencing (CITE-seq) analysis of Cas9-, RNP- and RNP/GSE56-electroporated human HSPCs. a-d**, Summary statistics for CITE-seq analysis, including total number of cells sequenced (a), mean number of genes per cell (b), mean number of transcripts per cell (Unique Molecular Identifiers, UMIs) (c) and mean mitochondrial read counts per cell (d) after quality control. **e**, Representative gene expression maps of lineage defining gene signatures, including HLF and MSI2 (hematopoietic stem cell [HSC] cluster 6); MPO (lymphoid-primed multipotent progenitors [LMPP] cluster 2 and granulocyte-monocyte progenitors [GMP] cluster 3); GATA2 (megakaryocytic-erythroid progenitors [MEP] cluster 5); and CD52 (common lymphoid progenitors [CLPs] cluster 1). **f**, UMAP visualization of HSC gene signatures extracted from Laurenti, et al. 2013; Eppert, et al. 2011; Fares, et al. 2017. **g**, Reconstruction of the hematopoietic hierarchy pseudo-time ordering by Slingshot. **h**, CITE-seq CD38, CD45RA, CD90 and CD49f expression within the identified clusters. Immunophenotypically defined LT-HSCs (CD38-CD45RA-CD90+CD49f+) were enriched in cluster 6. Associated with Fig. 4.

Extended Data Fig. 6

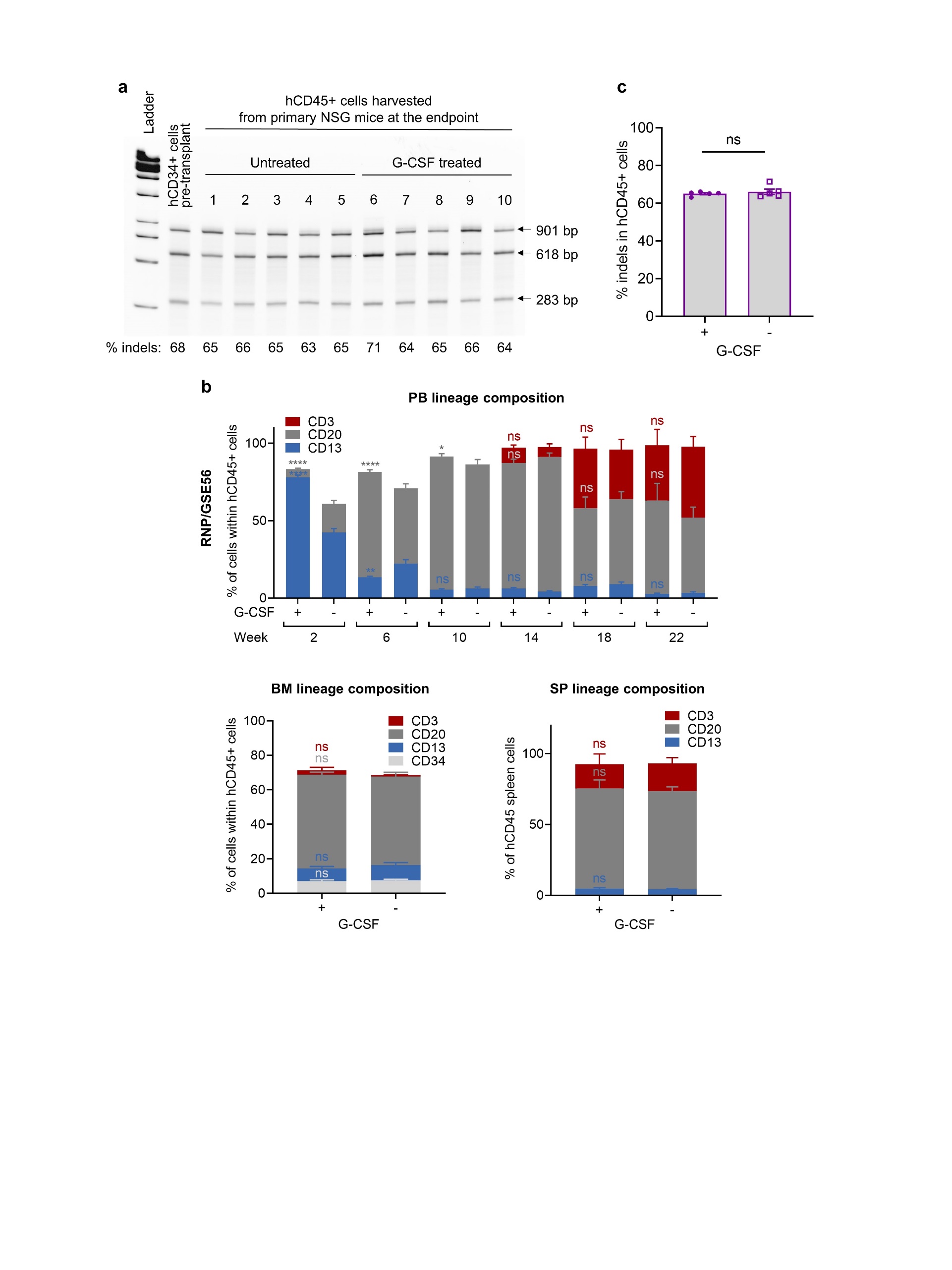

**Extended Data Fig. 6 | Analysis of gene editing frequency and lineage composition within human CD45+ cells after primary transplantation in the RNP/GSE56 group. a**, Gel image of the T7 endonuclease I (T7EI) assay for quantification of sgRNA/Cas9 RNP-mediated indels after electroporation (EP) in human CD34+ (hCD34+) cells pre-transplant or human CD45+ (hCD45+) cells isolated from murine bone marrow (BM) after primary transplantation. **b**, Lineage composition (CD3+ T lymphoid, CD20+ B lymphoid, CD13+ myeloid and CD34+ progenitor cells) within hCD45+ cells in the peripheral blood (PB), BM or spleen (SP) of NSG mice after primary transplantation in the RNP/GSE56 group. **c**, Summary of indel frequencies within hCD45+ cells harvested from the BM of NSG mice after primary transplantation as measured by T7EI assay. In panels **b and c**, data are displayed as mean ± standard error of the mean (SEM). Two-sided unpaired t-test was used. ns, not significant, ns, not significant, * p ≤ 0.05, ** p ≤ 0.01, **** p ≤ 0.0001. Associated with Fig. 6.
